# The molecular mechanism of positive allosteric modulation at the dopamine D1 receptor

**DOI:** 10.1101/2023.07.27.550907

**Authors:** Alexander Goldberg, Bing Xie, Lei Shi

## Abstract

The dopamine D1 receptor (D1R) is a promising target for treating various psychiatric disorders. While upregulation of D1R activity has shown potential in alleviating motor and cognitive symptoms, orthosteric agonists have limitations, restricting their clinical applications. However, the discovery of several allosteric compounds specifically targeting the D1R, such as LY3154207, has opened new therapeutic avenues. Based on the cryo-EM structures of the D1R, we conducted molecular dynamics simulations to investigate the binding and allosteric mechanisms of LY3154207. Our simulations revealed that LY3154207 preferred the horizontal orientation above intracellular loop 2 (IL2) and stabilized the helical conformation of IL2. Moreover, LY3154207 binding induced subtle yet significant changes in key structural motifs and their neighboring residues. Notably, a cluster of residues centered around the Na^+^ binding site became more compact, while interactions involving the PIF motif and its neighboring residues were loosened upon LY3154207 binding, consistent with their role in opening the intracellular crevice for receptor activation. Additionally, we identified an allosteric pathway likely responsible for the positive allosteric effect of LY3154207 in enhancing Gs protein coupling. This mechanistic understanding of LY3154207’s allosteric action at the D1R pave the way for the rational design of more potent and effective allosteric modulators.

## Introduction

G protein-coupled receptors (GPCRs) have been the focus of intensive efforts to discover and develop novel drugs for various diseases and disorders. These receptors are estimated to be the targets for nearly 40% of all prescribed drugs, underscoring their current importance and therapeutic potential [1, 2]. The dopamine D1 receptor (D1R), a class A rhodopsin-like GPCR, is one such receptor that has emerged as a promising target for the treatment of numerous psychiatric disorders [3], as it is known to play a key role in cognitive and locomotive processes [4, 5].

Despite the potential to treat Parkinson’s disease (PD) symptoms, it has been a persistent challenge to develop drugs that both selectively target the D1R and act at the central nervous system (CNS). Many D1R agonists are based on a catechol scaffold, which, however, possesses significant drawbacks that can impede their effectiveness. Specifically, catechol-based compounds tend to lack selectivity, exhibit poor bioavailability, and cannot penetrate the blood-brain barrier [6]. Currently, fenoldopam is the only FDA-approved D1R-selective agonist, and, as a poor CNS-penetrant, acts peripherally in treating hypertensive emergencies [7, 8].

In response to concerns surrounding the use of orthosteric drugs, significant effort has been directed towards developing allosteric compounds of GPCRs [3, 9]. Allosteric binding sites are typically located at distinct and divergent regions of the receptors, allowing allosteric drugs to avoid direct competition with orthosteric agonists as well as to exhibit significant selectivity [10]. Specifically, for the D1R, positive allosteric modulators (PAMs) may offer several advantages over orthosteric agonists, such as potentiating D1R activity with high selectivity or even cooperating with each other [11]. Additionally, D1R PAMs may be able to exert their allosteric effect without inducing tachyphylaxis, a side effect observed in some D1R agonists [12].

Several PAMs targeting the D1R at various sites have been discovered, synthesized, and pharmacologically characterized [11–15]. Among them are a group of compounds that share a similar structural scaffold containing a central tetrahydroisoquinoline (THIQ) ring, including 2-(2,6-dichlorophenyl)-1-[(1S,3R)-3-(hydroxymethyl)-5-(1-hydroxy-1-methyl-ethyl)-1-methyl-3,4-dihydro-1H-isoquinolin-2-yl]ethanone (DETQ), 2-(2,6-dichlorophenyl)-1-[(1S,3R)-3-(hydroxymethyl)-5-(2-hydroxy-2-methyl-propyl)-1-methyl-3,4-dihydro-1H-isoquinolin-2-yl]ethenone (DPTQ), and 2-(2,6-dichlorophenyl)-1-[(1S,3R)-3-(hydroxymethyl)-5-(3-hydroxy-3-methyl-butyl)-1-methyl-3,4-dihydro-1H-isoquinolin-2-yl]ethenone (LY3154207), of which DETQ and LY3154207 been found to efficiently cross the blood-brain barrier to act at the CNS [13]. These THIQ-based compounds may avoid some of the pharmacokinetic limitations of catechol-based drugs when targeting the D1R. DETQ has shown promise in preclinical studies and has been extensively studied in the context of the D1R allostery [12, 14]. LY3154207, which differs from DETQ only in the length of its extended two-carbon linker, demonstrates increased potency relative to DETQ [3, 13]. LY3154207 has progressed to clinical trials for the treatment of Lewy body dementia [13] and exhibited an acceptable safety and tolerability profile [16]. Additionally, clinical trials have shown that LY3154207 can improve motor symptoms related to Lewy body dementia, while either improving or not worsening non-motor symptoms associated with conventional dopaminergic treatment [17].

Recently, several high-resolution crystal and cryo-EM structures of the D1R in the active state have been published, providing crucial insights into the overall D1R structure, the mechanistic details of the coupling of D1R with Gs protein, and how D1R accommodates the agonists of various scaffolds and pharmacological properties [18–24]. Notably, the cryo-EM structures revealed that LY3154207 binds near intracellular loop 2 (IL2) at the receptor-membrane interface, interacting with residues in IL2, transmembrane segment 3 (TM3), and TM4 [19, 20, 22]. Furthermore, mutagenesis studies have indicated that IL2 is involved in the binding for both LY3154207 and DETQ [22, 25]. Interestingly, MLS1082, another D1R PAM that does not contain the THIQ moiety, has also been found to bind at IL2 [26, 27]. Interestingly, Cmpd-6FA and “Compound 1”, PAMs for the β2 adrenergic receptor (B2AR) and free fatty acid receptor 1, respectively, have been found to bind at similar pockets in the vicinity of IL2 [28, 29]. Therefore, the region encompassing IL2 emerges as a conserved allosteric site, suggesting the PAMs binding to this site may share some common elements in their allosteric mechanisms.

However, the D1R structures bound with and without LY3154207 show only limited differences, such as slightly deeper binding of the orthosteric ligand and only a small (<1.0 Å) global root-mean-square deviation (RMSD) between the D1R structures in the presence and absence of LY3154207 [19, 22]. While cryo-EM structures provide crucial structural information, an individual structure represents only a static snapshot of an ensemble of states and cannot provide information about the dynamics of the protein. Additionally, the experimental conditions necessary to acquire high-resolution cryo-EM structures of membrane proteins often do not mimic a physiological environment, and may mask functionally important effects, such as those of the PAMs bound between the protein-lipid interface. Furthermore, it is worth noting that structural changes may not be immediately apparent upon allosteric modulator binding [30]. As such, molecular dynamics (MD) simulations in a lipid environment provide an opportunity to explore a larger and more physiologically relevant conformational space to identify states that have not been resolved by microscopy or crystallography.

Thus, despite the promise demonstrated by LY3154207 and other D1R PAMs, the precise molecular mechanism underlying their allosteric modulations has not been fully characterized. An in-depth understanding the mechanism of action of LY3154207 may inform future drug discovery efforts, at not only the D1R but also other GPCRs. Therefore, we carried out a molecular modeling and dynamics simulation study of D1R in the presence and absence of LY3154207 with either dopamine or apomorphine bound in the orthosteric ligand binding pocket (OLBP). We identified structural and dynamic changes that may explain how LY3154207 enacts its allosteric effect on the D1R.

## Results and Discussion

To investigate the molecular mechanism of how PAM binding exerts its effect at the D1R, we conducted long-timescale atomistic molecular dynamics (MD) simulations in the presence and absence of LY3154207, with either dopamine or apomorphine bound in the orthosteric binding pocket (see Methods). We collected approximately 95 μs of MD simulation data in total for four conditions: D1R/dopamine-alone, D1R/dopamine-LY3154207, D1R/apomorphine-alone, and D1R/apomorphine-LY3154207 (see Table 1). We used these data to perform both quantitative and qualitative analyses of our systems to detect the differences induced by LY3154207 binding.

**Table 1.** Summary of MD simulations. Across all simulated conditions, a total of approximately 95 μs of simulation was collected.

| Ligand | Condition | LY3154207 orientation | Num traj | Length (μs) |
| --- | --- | --- | --- | --- |
| dopamine | D1R/dopamine-alone | - | 15 | 24.9 |
|  | D1R/dopamine-LY3154207 | horizontal | 9 | 21.3 |
|  |  | vertical | 3 | 3.6 |
| apomorphine | D1R/apomorphine-alone | - | 15 | 23.8 |
|  | D1R/apomorphine-LY3154207 | horizontal | 9 | 21.0 |

### LY3154207 is stable in the horizontal orientation

Despite the known location of the binding site of LY3154207, its binding orientation has not been conclusively determined. In the cryo-EM structure of the D1R bound with LY3154207 reported by Xiao et al, the PAM was found to bind “vertically”, oriented largely parallel to TMs 3 and 4 and perpendicular to IL2 [20]. Notably, prior to the first cryo-EM structure of the D1R, it was also predicted, based on an induced-fit docking work in a D1R homology model built from an active B2AR structure, that LY3154207 binds in a vertical orientation [13]. However, other D1R structures bound with LY3154207 indicate that LY3154207 may instead bind “horizontally” and lay parallel to IL2 [19, 22]. Remarkably, however, all groups that reported cryo-EM structures of the D1R bound with LY3154207 observed the presence of electron density corresponding to the “horizontal” orientation [19, 20, 22].

To address this controversy, we modeled and simulated systems with LY3154207 bound in either the “horizontal” or the “vertical” orientation observed in cryo-EM structures, in the presence of dopamine bound in the orthosteric ligand binding pocket (OLBP) (D1R/dopamine-LY3154207 (“horizontal”) and D1R/dopamine-LY3154207 (“vertical”), respectively, Figure 1A, D). Our results showed that the “horizontal” orientation was fairly stable, with the dichlorophenyl ring of LY3154207 sandwiched by an aromatic interaction with W123^3.52^ (superscripts denote Ballesteros-Weinstein residue indices [31]) and a cation-π interaction with R130^IL2^ (Figure 1B, Figure S1B). In contrast, we found that LY3154207 bound in the “vertical” orientation started to dissociate from the original binding pose within the first 200 ns of all the simulation trajectories and become considerably flexible and unstable. In one of the three trajectories of the D1R/dopamine-LY3154207 (“vertical”) condition, LY3154207 eventually dissociated from the allosteric binding site; in the other two trajectories, LY3154207 adopted transiently stable but divergent orientations that deviated drastically from the starting “vertical” orientation (Figure 1E).

**Figure 1.**
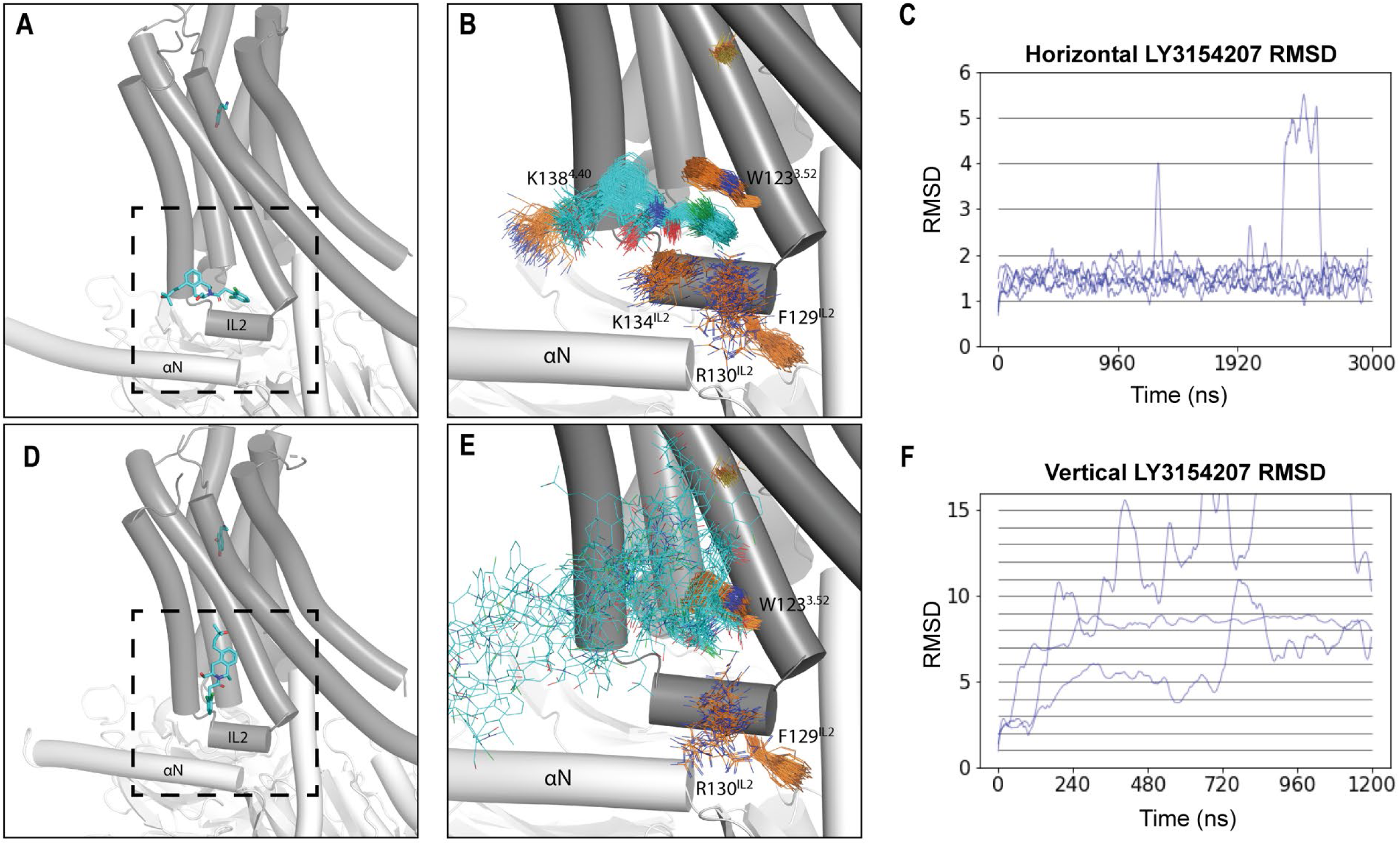
LY3154207 is stable in the horizontal orientation at its allosteric binding site of the D1R. Cryo-EM structures of the D1R (PDB 7LJD and 7CKZ) show that LY3154207 can be bound in the horizontal (A) and (D) vertical orientations, respectively. (B) Representative frames of the D1R/dopamine-LY3154207 (horizontal) condition show that LY3154207 is stable in the horizontal orientation. (C) RMSDs for the horizontal PAM binding orientation plateau around 1.5 Å in all D1R/dopamine-LY3154207 (horizontal) trajectories. (E) Representative frames of the D1R/dopamine-LY3154207 (vertical) condition reveal that LY3154207 was unstable in the vertical orientation. (F) RMSDs the vertical PAM binding orientation show that the PAM quickly dissociate from the binding pocket. The frames were aligned using TMs 3, 4, and IL2, while the evolution RMSD were calculated using the first frame of the trajectory as reference. LY3154207 is colored in cyan, the key binding site residues are in orange.

To quantitatively assess the stability of LY3154207 at the IL2 binding site, we calculated the ligand RMSD evolution along each trajectory. Our results revealed that the PAM was stable in the “horizontal” orientation in all the relevant trajectories, with RMSDs plateauing ∼1.5 Å along the evolution of the trajectories (Figures 1C and S1C). Furthermore, the pairwise RMSDs of LY3154207 were calculated using ten 2,000-frame random samples for each condition and resulted in an average pairwise RMSD of approximately 1.5 Å (Figure S2D). The similarity between the pairwise RMSD and the evolution RMSD suggests that the PAM remained relatively close to the initial pose during the simulations. In contrast, the evolution RMSD of the vertical LY3154207 sharply increased along the trajectories, which reflects an inability to persistently remain in this pose (Figure 1F).

For the “horizontal” orientation, upon visual inspection, we observed that the THIQ and dichlorophenyl moieties were much more stable than the 3-hydroxy-3-methylbutyl tail (Figures 1B and S2B, C, D). To confirm which parts of the PAM were most mobile, we calculated both the evolution and pairwise RMSD for three substructures of LY3154207 and found that the THIQ ring and dichlorophenyl moieties were indeed more stable than the 3-hydroxy-3-methylbutyl tail (Figure S2B, C, D).

Our finding that the “horizontal” orientation of LY3154207 is more stable than the “vertical” one is consistent with the proposal that the density for the reported vertical pose can be explained by the presence of a cholesterol molecule bound at this location [19]. The observation that mutating residues interacting with the vertical LY3154207 pose has a negative effect on cAMP accumulation [20, 25] may suggest that an interaction with cholesterol stabilizes LY3154207 and such mutations may disrupt this interaction [19]. Additionally, in the B2AR, the PAM Cmpd-6FA binds at IL2 in a “horizontal”-like pose, suggesting that PAMs may commonly adopt this orientation at the IL2-membrane interface (PDB: 6N48) [28]. For these reasons, we chose the “horizontal” binding pose for our following simulations and analyses.

### LY3154207 binding stabilizes intracellular loop 2 (IL2) in a helical conformation

It has been hypothesized that PAMs binding at IL2 stabilizes its helical structure, which in turn both reinforces the active state of the receptor and enhances G protein coupling [19, 20, 22, 28]. To assess whether LY3154207 indeed has a stabilizing effect on IL2, we analyzed the secondary structure for each residue of IL2 in each of our simulated conditions (see Methods). While residues 131-134 persistently adopted a helical conformation across all simulated conditions, we found that LY3154207 specifically stabilizes the helical conformation of residues 128-130 within IL2. In the D1R/dopamine-LY3154207 condition, we observed a significantly higher tendency for this portion of IL2 to be helical than in the D1R/dopamine-alone condition (Figure 2A, B). Upon conducting a thorough examination of our simulation results, we observed that the helical conformation of residue 128-130 still sporadically formed throughout the D1R/dopamine-alone simulations, albeit in a drastically lower frequency compared to the D1R/dopamine-LY3154207 simulations (Figure 2C, D). Notably, however, in both LY3154207 bound and unbound D1R cryo-EM structures [19], residues 128-130 are in essentially identical helical conformations (Figure S4). Consequently, we propose that the cryo-EM studies captured a stable state in the cryogenic temperature (<123 K), which may have suppressed the subtle yet critical dynamics of IL2, while our MD simulations conducted at 310 K can comprehensively characterize such dynamics and elucidate the impact of LY3154207.

**Figure 2.**
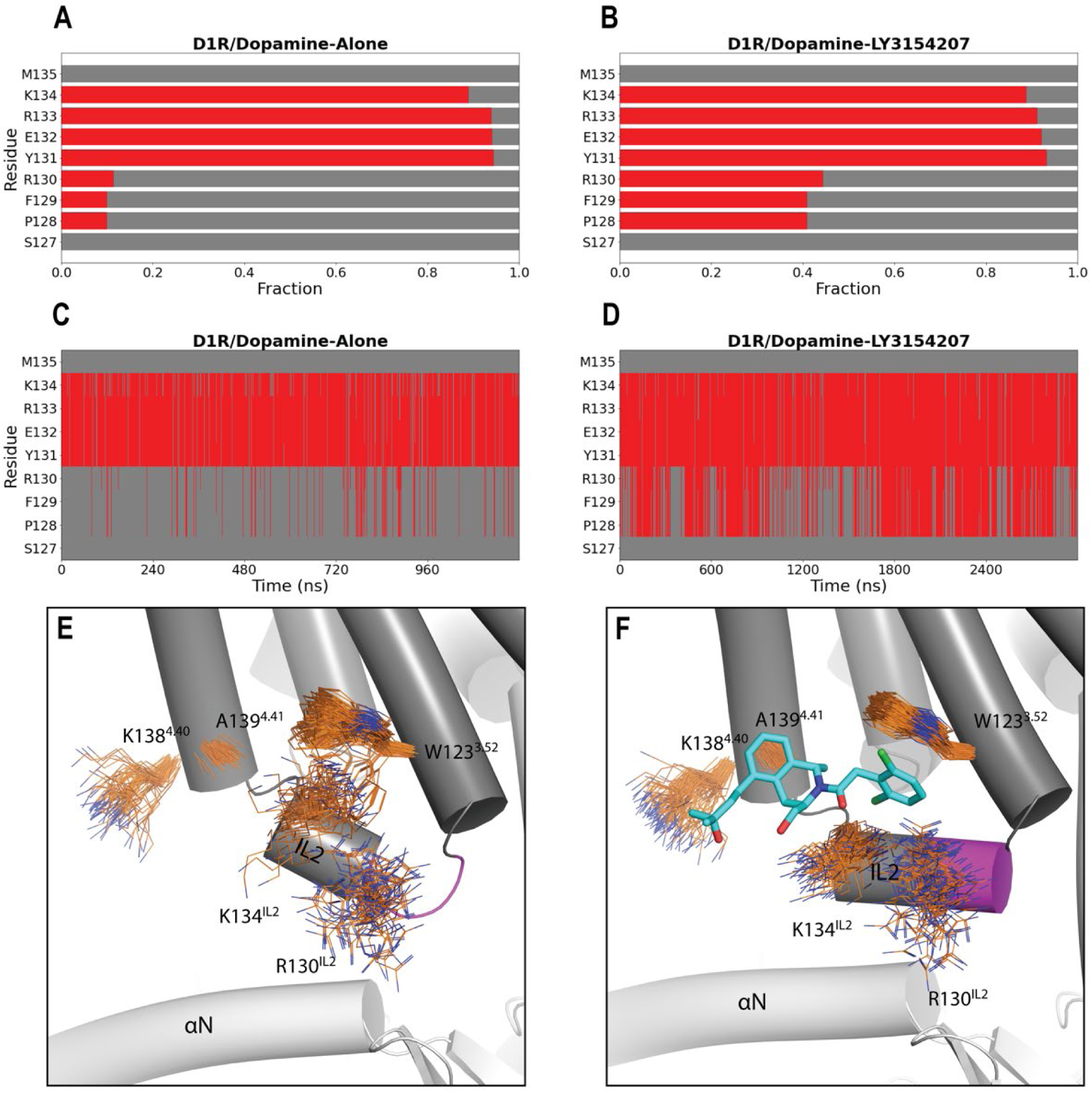
LY3154207 stabilizes the helical conformation for part of IL2. Comparing panels A and B, a greater propensity for residues P128, F129, and R130 of IL2 to adopt a helical conformation was observed when LY3154207 was bound (B) than when it was not (A). In panels C and D, the evolutions of secondary structure of IL2 are shown for the entire representative trajectories of the D1R/dopamine-alone and D1R/dopamine-LY3154207 conditions, respectively. Thus, in the D1R/dopamine-alone condition, this portion of IL2 could still be in a helical conformation throughout the trajectory but less frequently. In panels A-D, helical and loop conformation assignments are indicated with red and gray, respectively. In panels E and F, residues W123^3.52^, R130^IL2^, K134^IL2^, K138^4.40^, and A139^4.41^ contributing to the PAM binding are shown in line representation for the D1R/dopamine-alone (E) and PAM-bound D1R/dopamine-LY3154207 (F) conditions. The portion of IL2 including residues 128-130 is shown in magenta.

The helical conformation of residues 128-130 is partly attributed to the stabilizing interactions between the dichlorophenyl moiety of LY3154207 and W123^3.52^ and R130^IL2^, which form a sandwich-like configuration (Figure 2F). The functional roles of the key interactions formed between LY3154207 and W123^3.52^ and R130^IL2^ have been well documented by mutational studies in which altering these residues dramatically reduces the potency of the PAM as measured by cAMP accumulation [22, 25]. In addition, in the presence of LY3154207, there is a reduced flexibility of K134^IL2^ (Figure 2E, F). Despite K134^IL2^ having an α-helical characteristic in both the presence and absence of PAM, its polar interactions with LY3154207 restricted its movement. Consequently, IL2 adopts a closer and parallel alignment with the αN helix of the α subunit of Gs protein (Figure 2F). In the D1R/apomorphine system, we observed similar impacts of the PAM binding: when the PAM is bound, residues 128-130 are more frequently in helical conformation and IL2 was closer and more parallel to the αN helix as well (Figure S5).

Our simulation results also exhibited that A139^4.41^ forms a very stable hydrophobic interaction with the bulky THIQ ring. Interestingly, this is a divergent position between the two D1-like receptors, i.e., the D1R and the dopamine D5 receptor (D5R) [24]. Based on the high homology between these two receptors and the LY3154207-bound D1R structure, in the D5R, M^4.41^ is expected to clash with the bound LY3154207 [24]. Indeed, previous experimental results showed an approximately 1000-fold lower potency of DETQ at the D5R, manifesting high D1R-selectivity of these PAMs [25]. In a previous mutagenesis study, W123A, R130A, and A139L mutations of D1R all eliminated the PAM effect of LY3154207 [22]. As such, our results further validated that these residues are critical for the binding and allosteric effect of this PAM.

### LY3154207 has no strong impact on the orthosteric agonist binding

We then set out to identify the impact of LY3154207 binding on the rest of the D1R structure and first focused on the OLBP. While both the evolution and pairwise RMSD of LY3154207 were low (Figure 1), the orthosteric agonists bound in the OLBP had even lower RMSDs, indicating that the OLBP ligand is likely more stable. For all the simulated conditions, the average pairwise RMSD of the bound orthosteric agonists in the OLBP was between 0.7-1.2 Å (Figure S3).

MM/GBSA calculations revealed that there was no significant effect of LY3154207 binding on the binding free energy of dopamine (ΔG), with an average ΔΔG less than 1.4 kcal/mol between D1R/dopamine-alone and D1R/dopamine-LY3154207 (similar results were obtained in the comparison between the D1R/apomorphine-alone and D1R/ apomorphine-LY3154207 conditions) (Table S2). This suggests that the primary allosteric mechanism of LY3154207 is unlikely in inducing stronger binding of the bound orthosteric agonist.

### Network analysis identified significant impact of LY3154207 binding on the key structural motifs

We then carried out a network analysis to further investigate the structural and dynamic impact of LY3154207 binding on the D1R. This analysis involves transforming a protein structure, or a collection of similar structures (e.g., frames obtained from a MD simulation), into a graph comprising nodes and edges. Subsequently, graph theory is employed to examine and determine the mechanistic attributes of the protein. Specifically in this study, in each network, individual residues were represented as nodes, and each pair of nodes that had nonzero contact frequency in the corresponding simulated condition were connected with an edge (see Methods). With this approach, we calculated eigenvector centrality for each node. Eigenvector centrality serves as a metric for assessing the importance of a node to the network, where a higher eigenvector centrality score indicates that the corresponding node has a greater contribution to the network [32].

Strikingly, our network analysis results revealed that the nodes with the highest centrality scores across all simulated conditions predominantly consist of residues forming key structural motifs found in class A GPCRs, along with their neighboring residues. In particular, P206^5.50^, I111^3.40^, and F281^6.44^, which constitute the conserved PIF motif involved in receptor activation [33], and their neighboring residues, A109^3.38^, S110^3.39^, N113^3.42^, L114^3.43^, F203^5.47^, M210^5.54^, and W285^6.48^, were consistently observed among the top-scoring residues in all systems. Additionally, N327^7.49^, P328^7.50^, and Y331^7.53^ of the NPxxY motif [34] exhibited high scores in each of our systems. Notably, N327^7.49^ interacts with the Na^+^ binding residue D70^2.50^ [35–37]. Both D70^2.50^ and another Na^+^ binding residue S110^3.39^ scored high in all systems, as well as residues T37^1.46^, N41^1.50^, L66^2.46^, S69^2.49^, L72^2.52^, and V73^2.53^ in close vicinity (Tables 2 and S3). Furthermore, it has been proposed that the PIF and NPxxY motifs are connected through a linker leucine at position 3.43 in the B2AR, and their interactions play important roles in receptor activation and selective G protein/β-arrestin coupling [38]. Interestingly, L114^3.43^ was also among the highest scoring residues. These findings underscored the critical roles and positions of these structural motifs in the GPCR structures.

To evaluate the impact of the bound LY3154207, we calculated the differences between the scores in the presence and absence of LY3154207 for each top-scoring node in both the dopamine-bound and apomorphine-bound systems. Remarkably, our results showed a largely similar trend in the comparisons between D1R/dopamine-alone and D1R/dopamine-LY3154207 and between D1R/apomorphine-alone and D1R/apomorphine-LY3154207 (Tables 2 and S3). Specifically, LY3154207 binding resulted in higher scores for a cluster of nodes concentrated at the TMs 1, 2, 3, and 7 interface in both the D1R/dopamine and D1R/apomorphine systems (yellow residues in Figure 4 and S7, and Tables 2 and S3), compared to the corresponding conditions in the absence of LY3154207. This cluster includes nearly all aforementioned residues related to the Na^+^ binding motif, as well as P328^7.50^ from the NPxxY motif.

**Figure 3.**
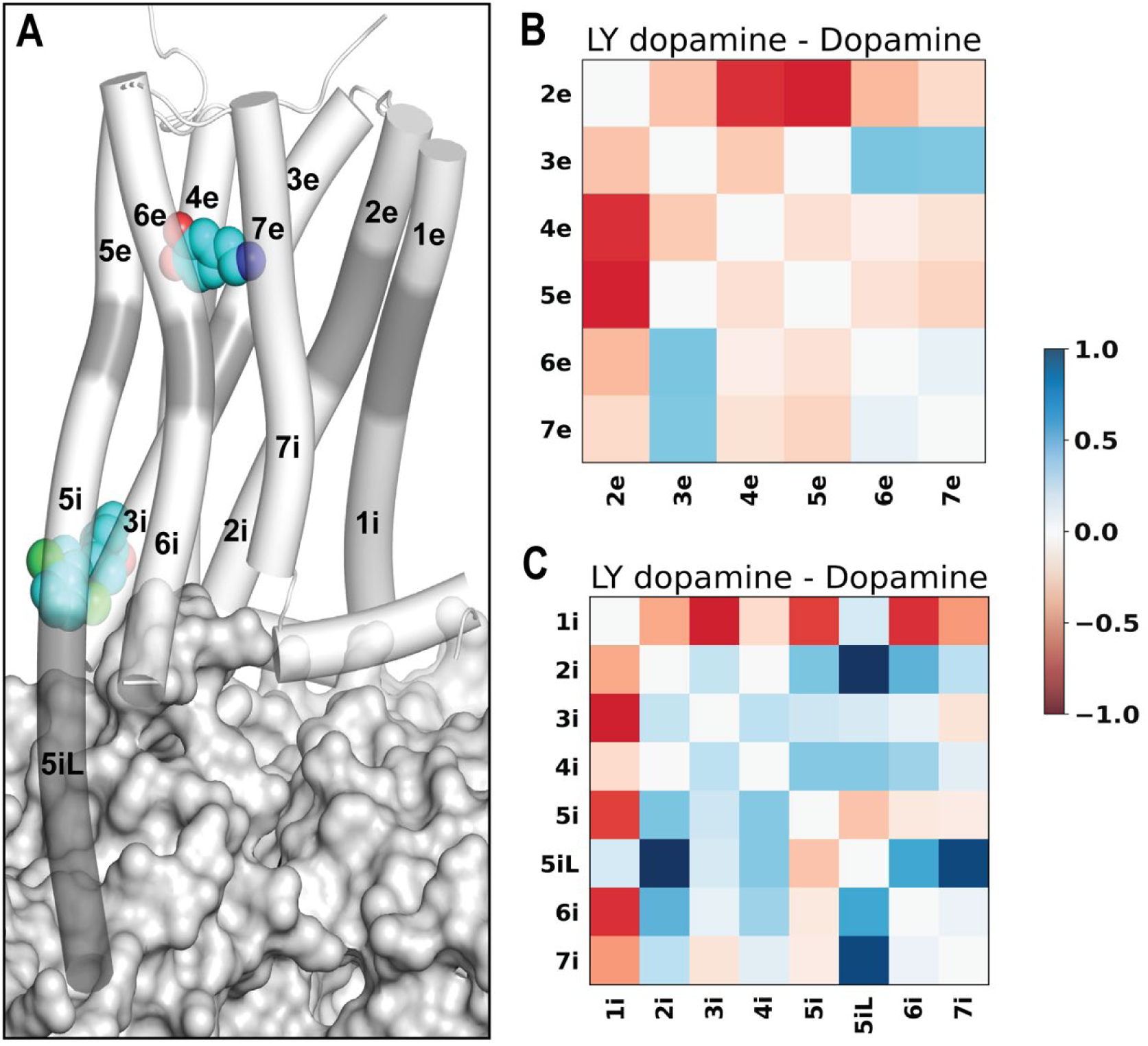
LY3154207 binding is associated with a tightening and loosening of the extracellular and intracellular regions of the receptor, respectively. (A) An overview of the D1R coupled with Gs protein and in complex with both dopamine and LY3154207 (PDB 7LJD). The extracellular and intracellular subsegments of the D1R are colored in white, while the middle segments are in gray (see Methods for their definitions). Our PIA analysis shows that the majority of extracellular subsegments have a tendency to move closer together (B), while the majority of intracellular subsegments have greater COM-COM distances (C) in the D1R/dopamine-LY3154207 compared to the D1R/dopamine-alone condition.

**Figure 4.**
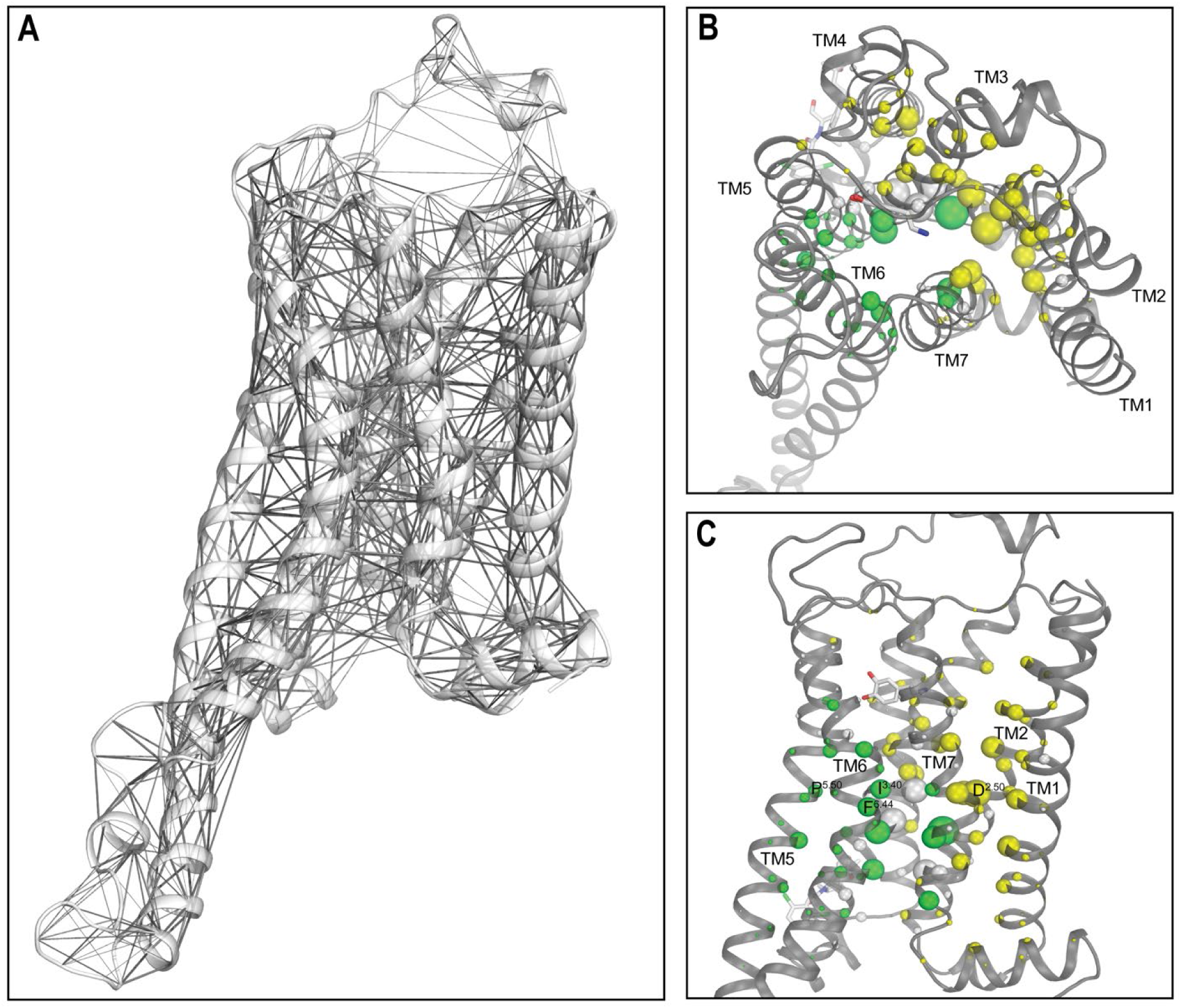
Network analysis identified impacts on several key structural motifs in the presence of LY3154207 in the D1R/dopamine systems. (A) A network connecting residues with larger than 0.2 contact frequency in the D1R/dopamine-LY3154207 condition. If the shortest heavy-atom distance between two residues was within 4.5 Å, we defined that they formed a contact. Edge radii are scaled by contact frequency. (B) Top and (C) side views of the D1R/dopamine-LY3154207 system with the residue nodes colored and scaled in radius according to the eigenvector centrality scores. All nodes with an absolute score difference exceeding 0.005 between the D1R/dopamine-LY3154207 and D1R/dopamine-alone conditions are highlighted in color. Thus, this eigenvector centrality analysis demonstrates that the presence of LY3154207 resulted in higher scores (yellow) for residues in the extracellular region enclosed by TMs 1, 2, 3, and 7, and lower scores (green) for residues in the intracellular region. Note that in addition to the TM residues discussed in text, S188^EL2^ has a significantly higher score in the LY3154207-bound D1R/dopamine-alone condition. Upon further inspection, we observed that this residue moved downward into the OLBP and interacts with the charged nitrogen of dopamine. However, this interaction was not observed D1R/apomorphine-LY3154207 condition since the bulky rings of apomorphine preclude such a movement of S188^EL2^.

The higher centrality scores of this cluster of residues are associated with higher contact frequencies with neighboring residues, and thereby a tightening of the TMs 1, 2, 3, and 7 interface. Such a tightening is consistent with the collapsing the Na^+^ binding site and the inward movement of the intracellular subsegment of TM7 (TM7i) near the NPxxY motif at this interface. The binding of a Na^+^ ion at the conserved Na^+^ binding site was associated with an inactive state of class A GPCRs [39], while the inward movement of TM7i coordinates with the outward movement of TM6i during receptor activation [40].

In contrast, a distinct cluster of residues comprising the PIF motif residues P206^5.50^, I111^3.40^, and F281^6.44^, and their neighboring L114^3.43^, F203^5.47^, and M210^5.54^, W285^6.48^, along with N327^7.49^ and Y331^7.53^ in the NPxxY motif, exhibited lower centrality scores in the presence of the bound PAM. These residues play critical roles in the conformational transition from the inactive to the active state. In particular, the outward movement of TM6i during receptor activation hinges around the helical turn encompassing positions F281^6.44^ and W285^6.48^ and is associated with the reconfigurations of both the PIF and NPxxY motifs. The lowered centrality scores of these residues are associated with less contact frequency with neighboring residues, which would facilitate the opening of the intracellular crevice of the receptor for coupling with the G protein.

It is noteworthy that both higher and lower centrality scores were observed for the residues within the NPxxY motif in the presence of the bound PAM. This can be attributed to the critical position of this motif between the two clusters of nodes centering around the Na^+^ binding and PIF motifs that demonstrated opposite trends in terms of centrality score changes. On one side, N327^7.49^ interacts with the Na^+^ binding D70^2.50^, while on the other side, Y331^7.53^ coordinates with Y214^5.58^ located below the PIF motif. In addition, N327^7.49^ and Y331^7.53^ face L114^3.43^, the sidechain of which is wedged between the PIF and NPxxY motifs and projects into the receptor core. Additionally, the configuration of the NPxxY motif is also associated with the reconfiguration of the 3_10_ helix in TM7 [41] which is located immediately above it and includes residues W321^7.43^, N323^7.45^, and S324^7.46^ identified among top-scoring nodes (Table 2).

**Table 2.** The network eigenvector centrality scores of the D1R/dopamine systems. Normalized eigenvector centrality scores for the D1R/dopamine systems in the presence and absence of LY3154207. Only residues with an eigenvector centrality score of 0.10 or greater in any of the conditions are shown, and the score difference and absolute score difference are colored (green to yellow and gray to white) according to their respective maximum and minimum values.

| Residue | BW index | D1R/DA | D1R/DA-LY | DA – DA-LY | abs diff |
| --- | --- | --- | --- | --- | --- |
| 37 | TM1.46 | 0.115 | 0.132 | -0.017 | 0.017 |
| 41 | TM1.50 | 0.118 | 0.140 | -0.022 | 0.022 |
| 62 | TM2.42 | 0.169 | 0.169 | 0.000 | 0.000 |
| 63 | TM2.43 | 0.114 | 0.111 | 0.003 | 0.003 |
| 65 | TM2.45 | 0.115 | 0.120 | -0.005 | 0.005 |
| 66 | TM2.46 | 0.217 | 0.204 | 0.013 | 0.013 |
| 67 | TM2.47 | 0.088 | 0.100 | -0.012 | 0.012 |
| 69 | TM2.49 | 0.162 | 0.168 | -0.006 | 0.006 |
| 70 | TM2.50 | 0.165 | 0.186 | -0.021 | 0.021 |
| 72 | TM2.52 | 0.096 | 0.105 | -0.009 | 0.009 |
| 73 | TM2.53 | 0.137 | 0.147 | -0.010 | 0.010 |
| 77 | TM2.57 | 0.097 | 0.107 | -0.010 | 0.010 |
| 78 | TM2.58 | 0.098 | 0.108 | -0.010 | 0.010 |
| 105 | TM3.34 | 0.093 | 0.106 | -0.013 | 0.013 |
| 106 | TM3.35 | 0.115 | 0.124 | -0.009 | 0.009 |
| 107 | TM3.36 | 0.082 | 0.083 | -0.001 | 0.001 |
| 109 | TM3.38 | 0.116 | 0.123 | -0.007 | 0.007 |
| 110 | TM3.39 | 0.172 | 0.172 | 0.000 | 0.000 |
| 111 | TM3.40 | 0.150 | 0.133 | 0.017 | 0.017 |
| 113 | TM3.42 | 0.174 | 0.177 | -0.003 | 0.003 |
| 114 | TM3.43 | 0.202 | 0.167 | 0.035 | 0.035 |
| 117 | TM3.46 | 0.163 | 0.146 | 0.017 | 0.017 |
| 118 | TM3.47 | 0.109 | 0.086 | 0.023 | 0.023 |
| 148 | TM4.50 | 0.135 | 0.144 | -0.009 | 0.009 |
| 151 | TM4.53 | 0.094 | 0.103 | -0.009 | 0.009 |
| 203 | TM5.47 | 0.111 | 0.094 | 0.017 | 0.017 |
| 206 | TM5.50 | 0.104 | 0.093 | 0.011 | 0.011 |
| 210 | TM5.54 | 0.142 | 0.116 | 0.026 | 0.026 |
| 214 | TM5.58 | 0.103 | 0.071 | 0.032 | 0.032 |
| 281 | TM6.44 | 0.151 | 0.125 | 0.026 | 0.026 |
| 285 | TM6.48 | 0.123 | 0.108 | 0.015 | 0.015 |
| 321 | TM7.43 | 0.117 | 0.126 | -0.009 | 0.009 |
| 323 | TM7.45 | 0.113 | 0.090 | 0.023 | 0.023 |
| 324 | TM7.46 | 0.128 | 0.142 | -0.014 | 0.014 |
| 327 | TM7.49 | 0.154 | 0.149 | 0.005 | 0.005 |
| 328 | TM7.50 | 0.090 | 0.115 | -0.025 | 0.025 |
| 331 | TM7.53 | 0.160 | 0.130 | 0.030 | 0.030 |

Interestingly, residues V63^2.43^, S69^2.49^, S107^3.36^, S110^3.39^ and N113^3.42^ occupying the central core of the receptor show high scores but little difference in the presence and absence of LY3154207. Indeed, residues which occupy dense regions of the receptor should be engaged in the interactions critical for the structural integrity. The lack of significant change in these residues suggests that these regions are not significantly impacted by PAM binding.

Taken together, our findings strongly indicate that the binding of LY3154207 reinforces the conformational changes that are closely associated with receptor activation.

### LY3154207 compacts and loosens the extracellular and intracellular regions of D1R, respectively

Next, we used the Protein Interaction Analyzer (PIA) [42, 43] to identify differences in the transmembrane (TM) domain of D1R between the PAM present and absent states. In this approach, we define subsegments of the TM domain, determine the location of their center of mass (COM), and measure COM-COM distances among the subsegments (see Methods). By comparing these distances between two conditions, we can detect structural effects that LY3154207 may have on specific regions of the TM domain.

The PIA results showed that in the comparison of the D1R/dopamine-LY3154207 and D1R/dopamine-alone conditions, a majority of extracellular subsegments interacted more closely with each other in the PAM bound condition. Specifically, with the exception of longer distances observed between the extracellular subsegment of TM3 (TM3e) and extracellular subsegments of TMs 6 and 7 (TM6e and TM7e, respectively), all the other pairwise distances of the extracellular subsegments decreased in the presence of the PAM (Figure 3B). This suggests that the PAM promotes tighter interactions among most extracellular subsegments, leading to a more compact arrangement. Furthermore, we observed noticeable lipid penetration near TM1e in the simulations when the PAM was not present. In contrast, we found no lipid penetration in our PAM-bound simulations, likely a result of tighter interactions in the extracellular region.

Our PIA analysis of the intracellular subsegments indicated a general loosening of the intracellular space in the presence of LY3154207 (Figures 3C and S6C), allowing for more extensive interactions between the intracellular crevice of the receptor and the G protein. In the D1R, TM5 is much longer than D2-like dopamine receptors, and the helix extends deeper into the intracellular milieu [24]. This extended conformation is also observed in other Gs coupled receptors [44]. To comprehensively analyze the effects on TM5, we divided the intracellular portion of the helix into two subsegments: TM5i, representing the intracellular portion of TM5 reaching the lipid boundary, and TM5iL, accounting for the extended helical portion of TM5 beyond the lipid boundary. Interestingly, TM5iL drifts away from the other intracellular subsegments in the D1R/dopamine-LY3154207, compared to the D1R/dopamine-alone condition (Figure 3C).

Intriguingly, even though in the network centrality analysis, we observed a highly similar trend in the comparisons between the two D1R/dopamine conditions and between the two D1R/apomorphine conditions, our PIA analysis revealed limited commonality in specific distance changes, particularly on the extracellular side (compare Figures 3 and S6). This discrepancy can be attributed to the distinct impacts of different agonist bindings, which may potentially overshadow certain conformational changes associated with receptor activation when assessed through individual distances. Nevertheless, it is important to note that in the presence of the bound PAM, in both the D1R/dopamine and D1R/apomorphine systems, the extracellular subsegments exhibited increased compactness, while the intracellular subsegments were loosened (Figures 3 and S6).

### A potential allosteric pathway at the receptor-G protein interface

To better understand the impact of LY3154207 on the interactions between the D1R and Gα, we performed contact frequency difference analysis (see Methods). Our results showed that the downward movement of IL2 resulting from the binding of LY3154207 allowed IL2 to form more interactions with the G protein. Specifically, R133^IL2^ of D1R has increased contact frequencies with A39, K216, and V217 of Gα (Figures 5B, C and S9A, B), because the bound LY3154207 restricted the rotational freedom of R133^IL2^, causing it to be constrained to interact more closely with Gα. Related to this difference, we found that the interactions between Y131^IL2^ of D1R and H387 and Y391 in the last two turns of the α5 helix of Gα were weakened in both the PAM-bound D1R/dopamine-LY3154207 and D1R/apomorphine-LY3154207 conditions (Figures 5B, C and S9A, B).

**Figure 5.**
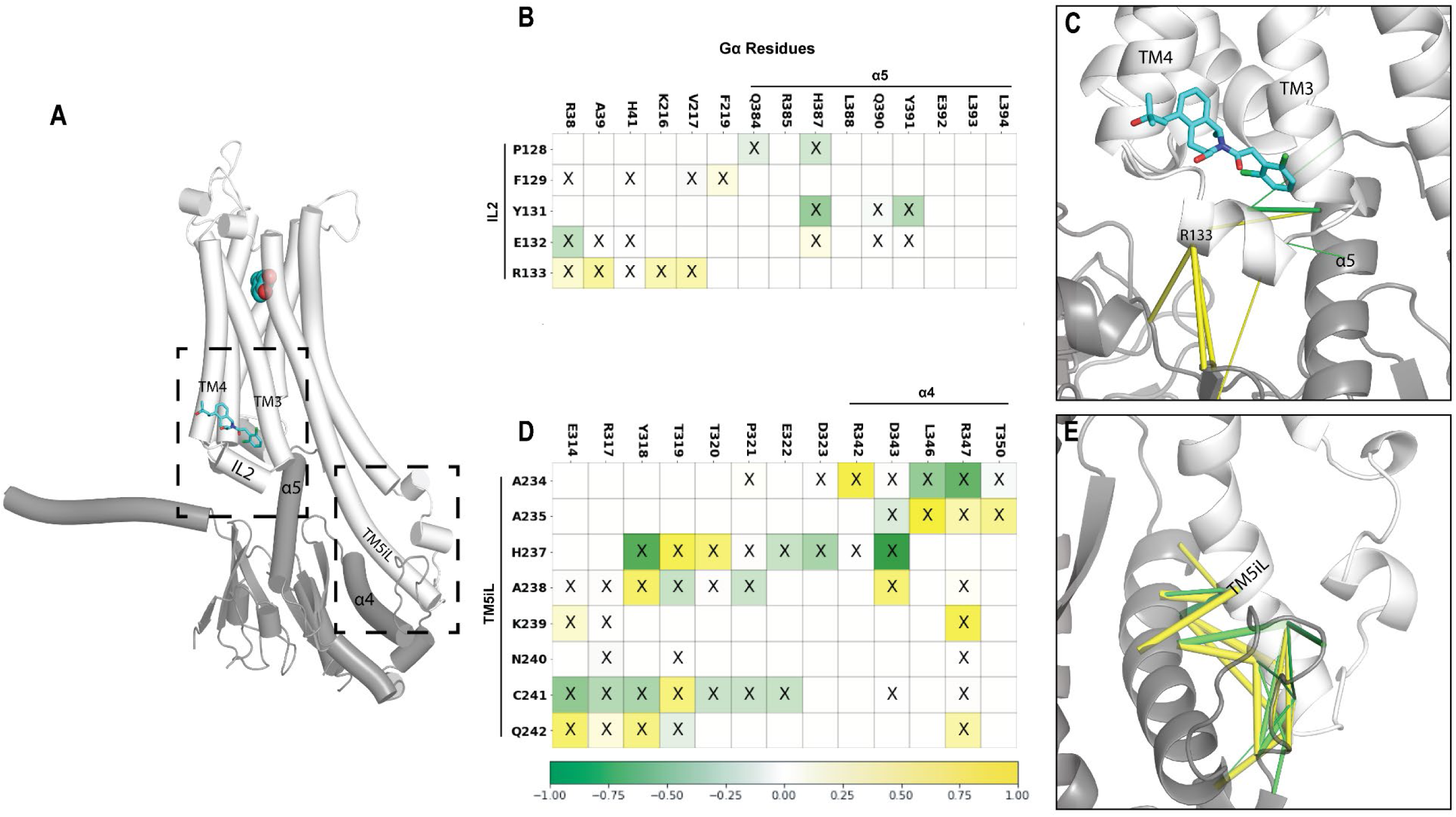
Contact frequency analysis between the D1R and Gα_s_ reveals an allosteric pathway in the D1R/dopamine systems. (A) Overview of the D1R/dopamine-LY3154207 condition. Dashed boxes indicate the regions shown in panels C and E. (B) The contact map for the interface between D1R/IL2 and Gα residues. Cells containing an ‘X’ represent the corresponding residue pairs forming a contact in both the D1R/dopamine-LY3154207 and D1R/dopamine-alone conditions. More frequent interactions in the presence of PAM are colored in yellow, and less frequent interactions are in green. These divergent interactions are mapped on the D1R/Gα model shown in panel C. (C) In the presence of PAM, IL2 makes more consistent interactions with αN of Gα, specifically R133^IL2^. For comparison of the same two conditions, panel D shows the contact map for the interface between D1R/TM5iL and Gα residues, while the divergent interactions are mapped to the model in panel (E). All contacts with a frequency difference greater than or equal to 0.10 are shown, and tubes are scaled to the frequency difference.

In the D1R/dopamine-LY3154207 condition, this impact of PAM binding propagated to the extended portion of TM5 of D1R, i.e., TM5iL, resulting in its distant portion (residues 234^5.78^ to 242^5.86^) to engage in more frequent contacts with R342 to T350 in the α4 helix of Gα (Figure 5D, E). This change involved a slight rotation of TM5iL, as one residue was oriented closer to Gα while its immediate neighbor rotated away from Gα. For example, in the presence of the bound PAM, D1R-A238 has higher contact frequency with Gα-Y318, while D1R-H237 has a tendency to move away from Gα-Y318 (Figure 5D). A similar trend of TM5iL rearrangement was also found in the comparison of D1R/apomorphine-LY3154207 and D1R/apomorphine-alone conditions (Figure S9C, D).

In addition, by comparing the two PAM-bound conditions, we found that IL3, which extends from TM5iL, has a relatively convergent and more structured conformation (Figure S8C, D), whereas the IL3 conformation in the dopamine- and apomorphine-alone conditions are fairly divergent and less structured (Figure S8A, B). It is likely that this convergent conformation observed in the presence of LY3154207 is related the slight rotation of TM5iL described above.

Taken together, we identified the converged structural changes at the receptor-G protein interface in both pairs of the simulated conditions. These changes led to the formation of an allosteric pathway, originating from the binding of the PAM near the IL2 binding pocket and extending to TM5iL and IL3. Notably, TM5iL, which is helical in nature, has been identified as a unique feature of Gs-coupled receptors [44]. Hence, our findings unveil intriguing properties of the PAM allosteric mechanism that may be specific to the Gs-coupled receptors.

## Conclusion

In this study, we carried out long-timescale MD simulations of the D1R-Gs complex, both in the presence and absence LY3154207, a D1R-selective PAM. Our aim was to identify and characterize structural and dynamic changes induced by the binding of LY3154207 to the D1R. Through network analysis of the simulation results, we found that key structural motifs conserved in class A GPCRs and their neighboring residues had the highest centrality scores across all simulated conditions. Among these residues, LY3154207 binding increased centrality scores in a cluster of residues centered at Na^+^ binding residues, tightening the interface among TMs 1, 2, and 7. In contrast, the PIF motif and neighboring residues exhibited lower centrality scores in the presence of the bound PAM, consistent with their involvement in opening of the intracellular crevice associated with receptor activation. These trends are also in line with the compaction of the extracellular region and an opening of the intracellular region of the receptor revealed by our PIA analysis. Importantly, we observed convergent structural changes at the interface between the D1R and the Gα in both pairs of simulated conditions. These changes revealed an allosteric pathway that originated from the binding of the PAM near the IL2 binding pocket and extended towards TM5iL and IL3. Interestingly, LY3154207 had no significant impact on the OLBP, leading us to propose that its main allosteric effect is to enhance G protein coupling. In summary, our findings shed light on the structural dynamics of D1R upon LY3154207 binding and provide insights into the allosteric modulation of G protein coupling by this D1R-selective PAM.

## Methods

### Homology modelling

Homology modeling for the D1R in complex with Gs was carried out with Modeller (version 9.24) [45], using the cryo-EM structures of the D1R bound with dopamine and LY3154207 (PDB: 7LJD) [22] or with dopamine alone (PDB: 7F1Z) [46] as the templates for both the D1R/dopamine-alone and D1R/dopamine-LY3154207 conditions. Specifically, 100 models were generated and assessed by the DOPE score. A model with a low DOPE score while having the loops in feasible orientations with respect to the membrane, was selected for further refinement of the loop regions without any template. 100 models were built by Modeller for each of two loop refinements, one for the extracellular loops and one for the intracellular loops. A low DOPE score model with feasible loop orientations was selected from each stage to use as input in the next stage. IL3 (residues 242-264) was further refined using the next-generation kinematic closure (NGK) protocol implemented in Rosetta (version 2019.47.61047) [47], which considers the context of conformational sampling of local regions. 2,000 models were generated, and the resulting low energy model with feasible IL3 orientation was selected from a dominant cluster that shared overlapping α helical character with the base of TM5. Cys347 in H8 was palmitoylated using CHARMM-GUI (version 3.7) [48].

The apomorphine systems were built by modifying equilibrated frames from the dopamine models, and apomorphine was grafted into the OLBP in place of dopamine using a published structure of the D1R bound with apomorphine as reference (PDB: 7JVQ) [21].

### Molecular dynamics simulations

The D1R models were further processed in Maestro (Schrodinger version 2021-3) using the Protein Preparation Wizard. D70^2.50^ and D120^3.49^ were protonated to their neutral forms as assumed in the active state of aminergic GPCRs [49] and the nitrogen of dopamine was protonated to a positively charged state. The D1R/dopamine simulation systems were built using the Desmond System Builder (Schrodinger version 2021-3). Briefly, the models were immersed in explicit 1-palmitoyl-2-oleoyl-*sn*-glycero-3-phosphocholine (POPC) lipid bilayer. The simple point charge (SPC) water model was used to solvate the system, the net charge of the system was neutralized by Cl^-^ ions, and then 0.15 M NaCl was added. The resulting system consists of approximately 182,000 atoms. The equilibrated system has approximate dimensions of 108×105×161 Å^3^. Histidine protonation states were identical in all models and are listed in Table S1.

Desmond MD systems (D. E. Shaw Research, New York, NY) was used for the MD simulations using the OPLS4 force field [50]. The force field parameters for LY3154207 and apomorphine were further optimized quantum mechanically by the force field builder of the Schrodinger Suites (version 2021-3). Similar to our previous simulation protocols used for GPCRs [36, 51, 52], the system was initially minimized and equilibrated with restraints on the ligand heavy atoms and protein backbone atoms with a force constant of 1.0 kcal/mol/Å. The NPγT ensemble was used with constant temperature and pressure maintained with Langevin dynamics. Specifically, 1 atm constant pressure was achieved with the hybrid Nose-Hoover Langevin piston method on an anisotropic flexible periodic cell with a constant surface tension (x-y plane). Restraints on the systems were gradually released over approximately 250 ns to prepare them for long-timescale simulations. The restraints on the D1R loops, TM1, the D1R TM domain, and the bound ligands were released in this order. In the production runs at 310 K, all restraints of the D1R were released while most of the α carbons restraints of G protein were retained, except for the Gα residues 303-351 (α4-αG loop and α4 helix) and 383-394 (α5 helix). For each condition, we collected at least three trajectories starting from different random number seeds. Overall, 51 trajectories, with an aggregated simulated time of approximately 95 μs, were collected (Table 1).

### DSSP Structure prediction

The secondary structure assignment to three broad categories: helix, sheet, and coil, was carried out with MDtraj (version 1.9.4) [53] using the Define Secondary Structure of Proteins (DSSP) algorithm [54]. The assignment is primarily based on the electrostatic interaction energy calculated for backbone atoms [54]. The assignments were calculated for each residue of IL2 in each simulated condition. Calculations were performed on a 20,000-frame bootstrap sampling of all representative trajectories for each system.

### Ligand RMSD calculation

In calculating the ligand evolution RMSD, we first aligned the ligand binding pocket residues of each MD frame to those of the first frame of the corresponding MD trajectory, then computed the RMSD of the heavy atoms of the ligand. Alignment for the OLBP ligand calculations was performed using Cα of the residues within 4.5 Å of the ligand in any condition. Alignment for the PAM RMSD calculations were performed using Cα of TM3, 4 and IL2 residues (112^3.41^-150^4.52^). The pairwise RMSD was calculated using the same alignment scheme with a dataset of ten 2,000-frame random samples for each condition. For each sample, we calculate the pairwise RMSDs for all possible pairs, and then averaged the resulting RMSD. The reported average pairwise RMSD is the average of ten random sample averages and the standard deviation represents the deviation across those ten averages.

### MM/GBSA Calculation

Binding free energies between the bound ligands and the D1R were estimated with the Molecular mechanics/generalized Born surface area calculations (MM/GBSA) method, using the same force field in the MD simulations for the proteins and ligands, but with VSGB2.1 solvation model [55]. We extracted frames every 3 ns from the production runs to carry out the MM/GBSA calculations using the thermal_mmgbsa.py script from the Schrodinger suite (version 2021-3). The binding free energies for each condition were the averages of the selected frames.

### Protein Interaction Analyzer (PIA)

An average COM-COM distance calculation was performed for each bootstrapped trajectory. The following residue ranges were used to define transmembrane subsegments for all PIA calculations: 1e (21^1.30^-27^1.36^), 1m (28^1.37^-36^1.45^), 1i (37^1.46^-50^1.59^), 2i (58^2.38^-71^2.51^), 2m (72^2.52^-80^2.60^), 2e (81^2.61^-86^2.66^), 3e (93^3.22^-106^3.35^), 3m (107^3.36^-111^3.40^), 3i (112^3.41^-126^3.55^), 4i (137^4.39^-147^4.49^), 4m (148^4.50^-153^4.55^), 4e (154^4.56^-160^4.62^), 5e (192^5.36^-201^5.45^), 5m (202^5.46^-206^5.50^), 5i (207^5.51^-219^5.63^), 5iL (220^5.64^-242^5.86^), 6i (267^6.30^-280^6.43^), 6m (281^6.44^-285^6.48^), 6e (286^6.49^-297^6.60^), 7e (310^7.32^-321^7.43^), and 7i (322^7.44^-332^7.54^). The resulting distances were compared between systems and the differences were plotted for each subsegment COM-COM distance. Calculations were performed on a 20,000-frame bootstrap sampling of all representative trajectories for each system.

### Eigenvector centrality

In network analysis of protein structures, residues are commonly reduced to individual nodes. Two nodes can then be connected by an edge if their distance is below a certain threshold, which is defined as a contact. In this study, contacts (heavy-atom distance <= 4.5 Å) were identified among all residues of the receptor for a given MD frame. Using ten 2,000-frame bootstrapping samples for each condition, we calculated the frequency for each contact and built average contact frequency matrix. The contact frequency matrix was converted to an adjacency matrix, where nonzero frequency resulted in an edge connecting any pair of nodes separated by two or more residues in sequence. A network was built from this adjacency matrix using igraph (version 0.10.1) and the eigenvector centrality score of each node was calculated with edge weights equal to the contact frequencies between each pair of contacting residues. Note that the eigenvector centrality score is proportional to the sum of centrality scores of its neighbors, and so each node score is influenced by the score of nodes to which it is connected.

### Contact frequency difference analysis of the D1R-Gα interface

In an MD frame, if the shortest heavy-atom distance between a D1R residue and a G residue was within 4.5 Å, we defined that they formed a contact. For each simulated condition, we first determined the frequency for each contact in a 2,000-frame bootstrap sampling. Subsequently, we calculated the average frequency across ten such samplings. Finally, we compared the contact frequencies between the conditions where the PAM was bound and unbound by computing the differences of the corresponding contacts.

## Author Contribution

A.G. and L.S. designed the study. A.G. and B.X. carried out computational modeling, simulations, and analysis. All authors took part in interpreting the results. A.G. prepared and wrote the initial draft, all authors finalized manuscript.

## Declarations of Competing Interests

No potential conflict of interest was reported by all authors.

## Acknowledgements

We thank David Sibley for insightful discussions. Support for this research was provided by the National Institute on Drug Abuse–Intramural Research Program, Z1A DA000606 (L.S.). This work utilized the computational resources of the NIH HPC Biowulf cluster (http://hpc.nih.gov).

## Supplementary Information

**Figure S1.**
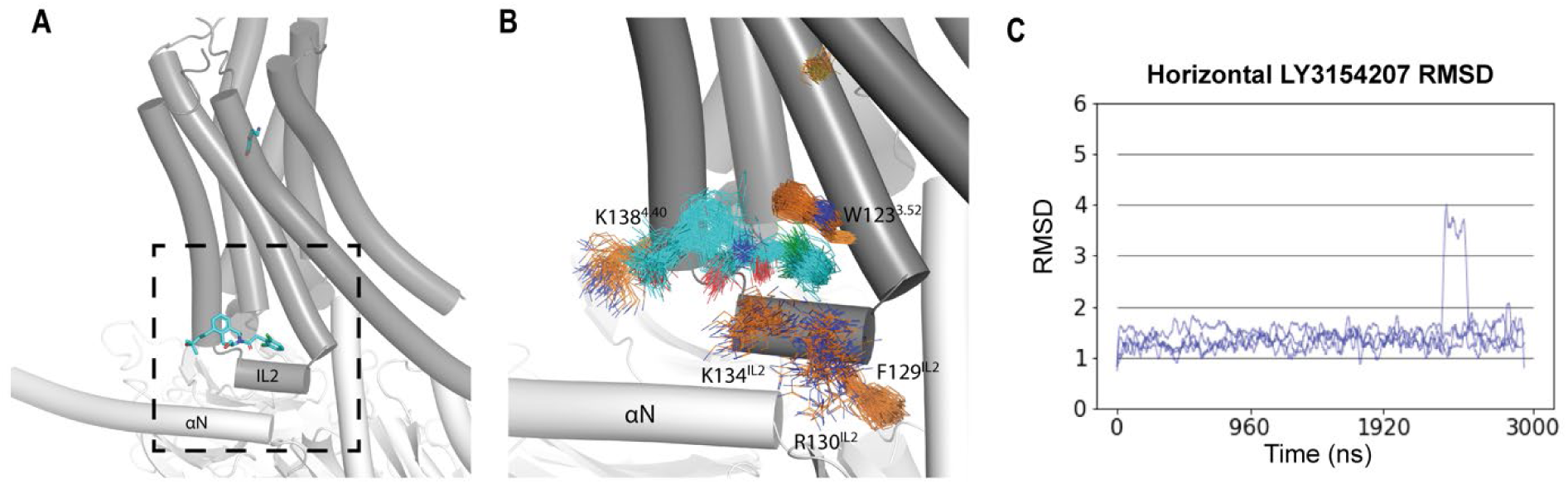
Stability of LY3154207 at IL2 in the D1R/apomorphine systems. Cryo-EM structures of LY3154207 bound in the (A) horizontal orientation (PDB 7LJD). (B) Representative frames of the D1R/apomorphine-LY3154207 (horizontal) condition show that LY3154207 is stable in the horizontal orientation. (C) RMSDs for the horizontal PAM binding orientation plateau around 1.5 Å in all trajectories.

**Figure S2.**
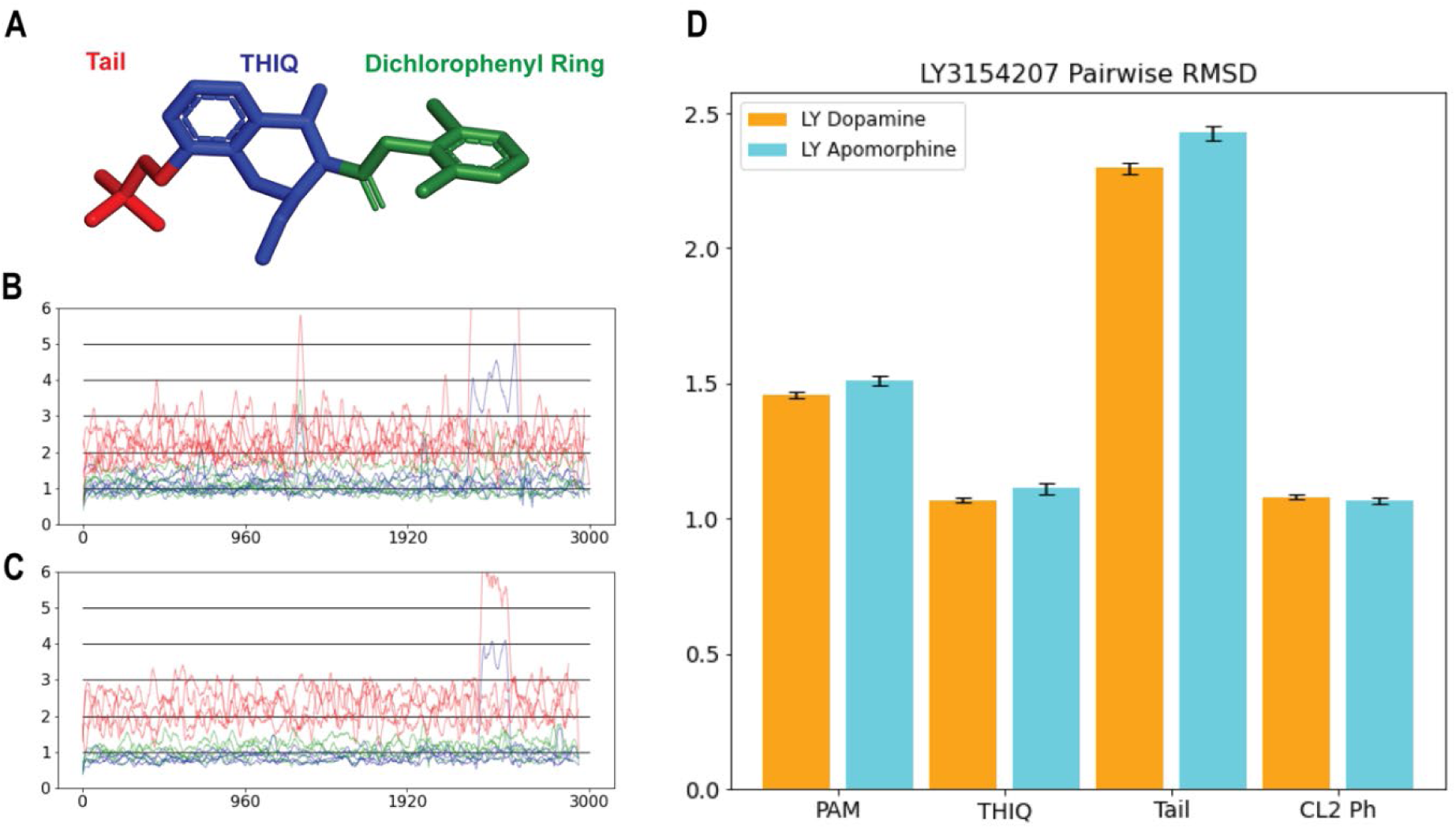
Stability of LY3154207 at IL2 in all simulated systems. (A) The LY3154207 is divided into three substructures. RMSDs for the 3-hydroxy-3-methylbutyl tail (red), THIQ (blue), and dichlorophenyl ring (green) are plotted for both the D1R/dopamine-LY3154207 (B) and D1R/apomorphine-LY3154207 (C) conditions. (D) Pairwise RMSDs calculated as the average of ten 2000-frame random sample averages demonstrate considerable stability of the THIQ and dichlorophenyl ring moieties. Error bars represent the standard deviation across the 10 average pairwise RMSDs.

**Figure S3.**
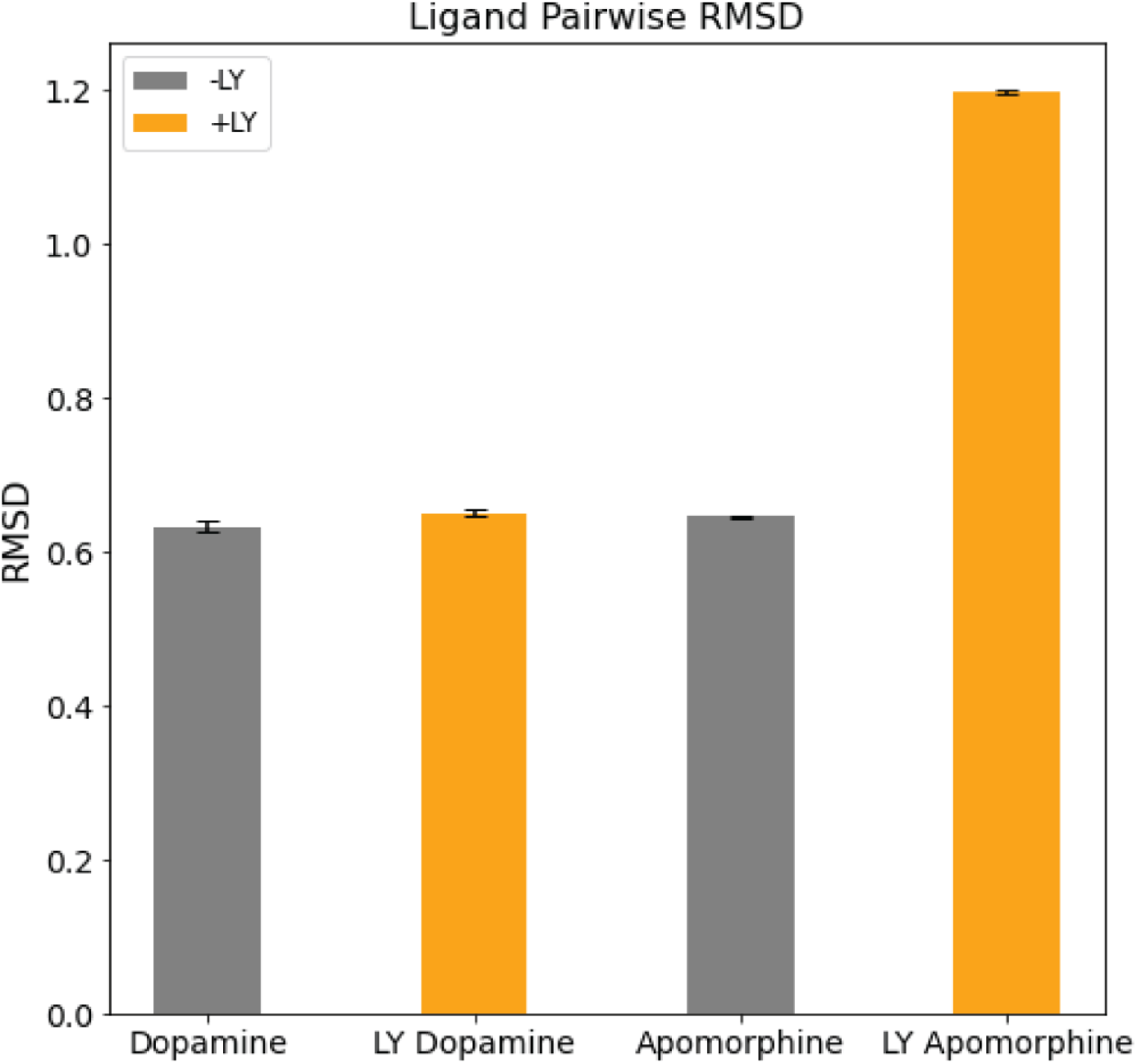
Pairwise RMSD for ligands bound in the OLBP in the presence and absence of LY3154207. The ligand in the OLBP is stable with pairwise RMSDs below 1.2 Å. PAM binding has no significant effect on the stability of dopamine in the OLBP, however there is an increase in the pairwise RMSD for apomorphine in the D1R/apomorphine-LY3154207 condition. Each bar represents the average of average pairwise RMSDs from ten 2000-frame random samples, and error bars represent the standard deviation across the ten averages.

**Figure S4.**
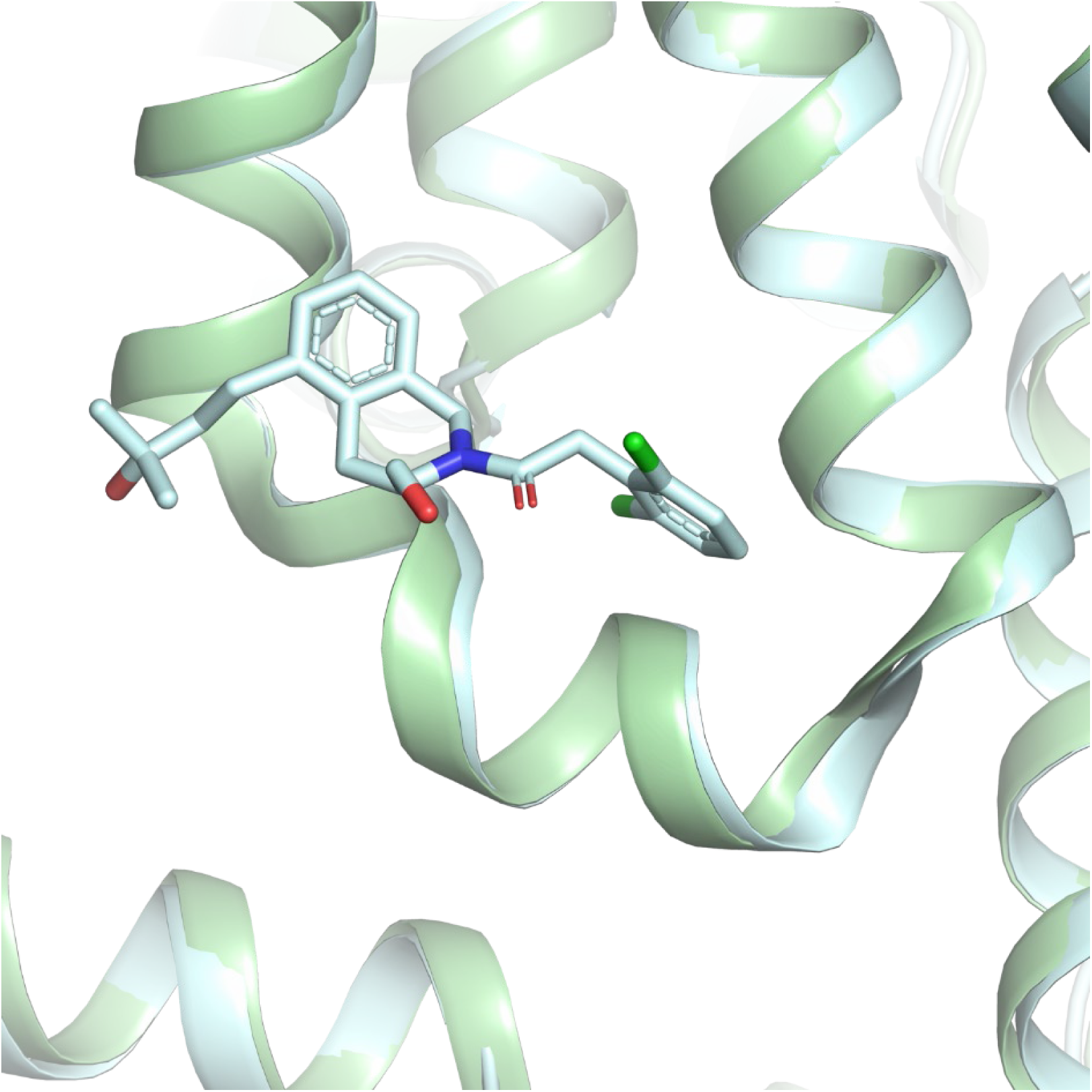
Recent cryo-EM structures of D1R/dopamine in the presence and absence of LY3154207 show nearly identical IL2 conformations. A comparison between two recently published cryo-EM structures of the D1R/dopamine system aligned by the Cα atoms of the receptor residues in the presence (PDB 7X2F, shown in pale cyan) and absence (PDB 7F0T, shown in pale green) of LY3154207. Both structures are in the absence of GDP but in complex with Nb35.

**Figure S5.**
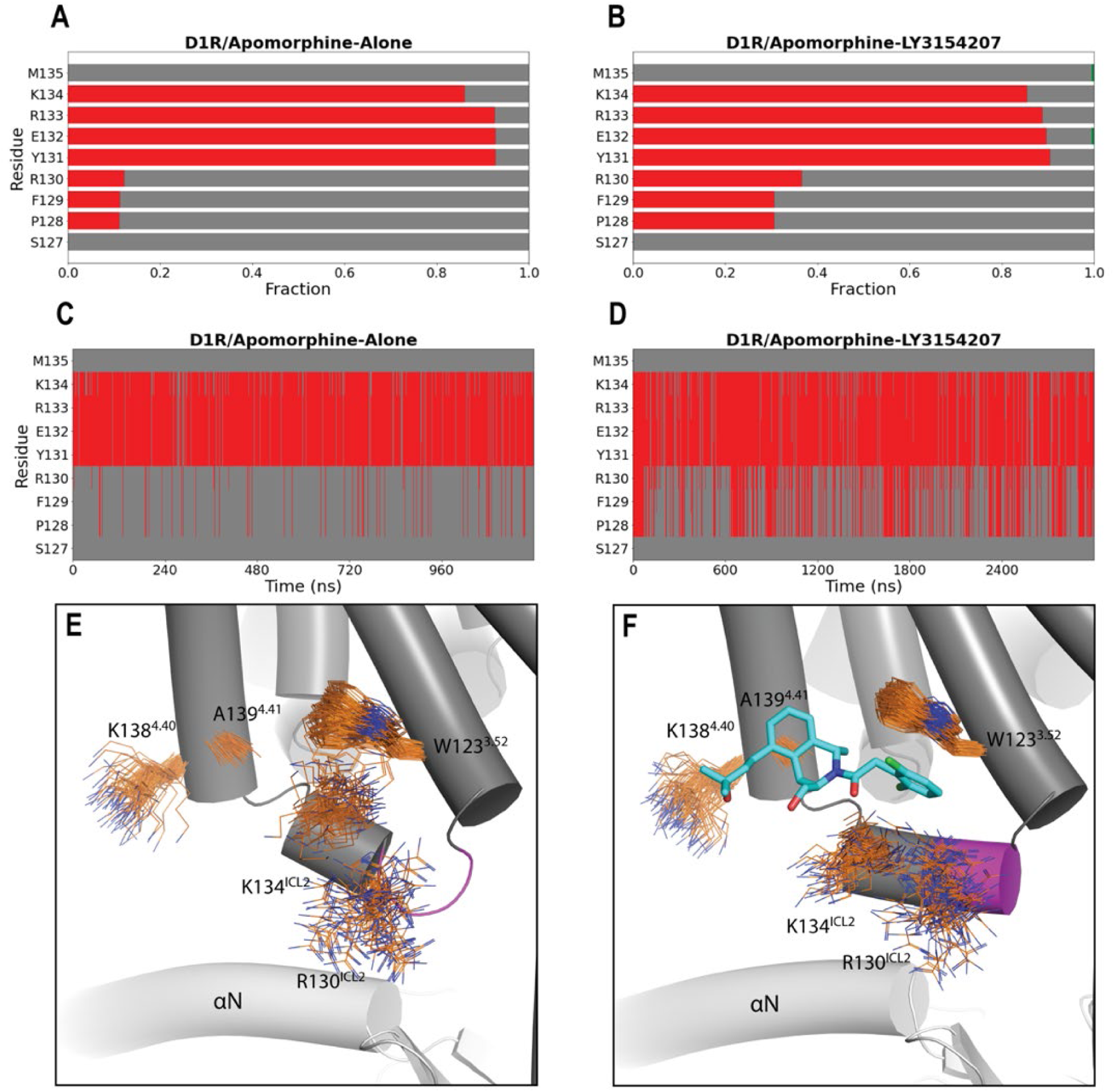
LY3154207 stabilizes a helical structure for intracellular loop 2 in the D1R/apomorphine system. There is a greater helical frequency for residues P128, F129, and R130 when LY3154207 is bound (B) versus when it is not (A). In panels C and D, the evolutions of secondary structure of IL2 are shown for the entire representative trajectories of the D1R/apomorphine-alone and D1R/apomorphine-LY3154207 conditions, respectively. In the absence of PAM, this portion of IL2 could still adopt a helical conformation throughout the trajectory but less frequently. In panels A-D, helical and loop conformation assignments are indicated with red and gray, respectively. In panels E and F, residues W123^3.52^, R130^IL2^, K134^IL2^, K138^4.40^, and A139^4.41^, which contribute to PAM binding, are shown in line representation for the D1R/apomorphine-alone (E) and D1R/apomorphine-LY3154207 (F) conditions. The portion of IL2 including residues 128-130 is highlighted in magenta.

**Figure S6.**
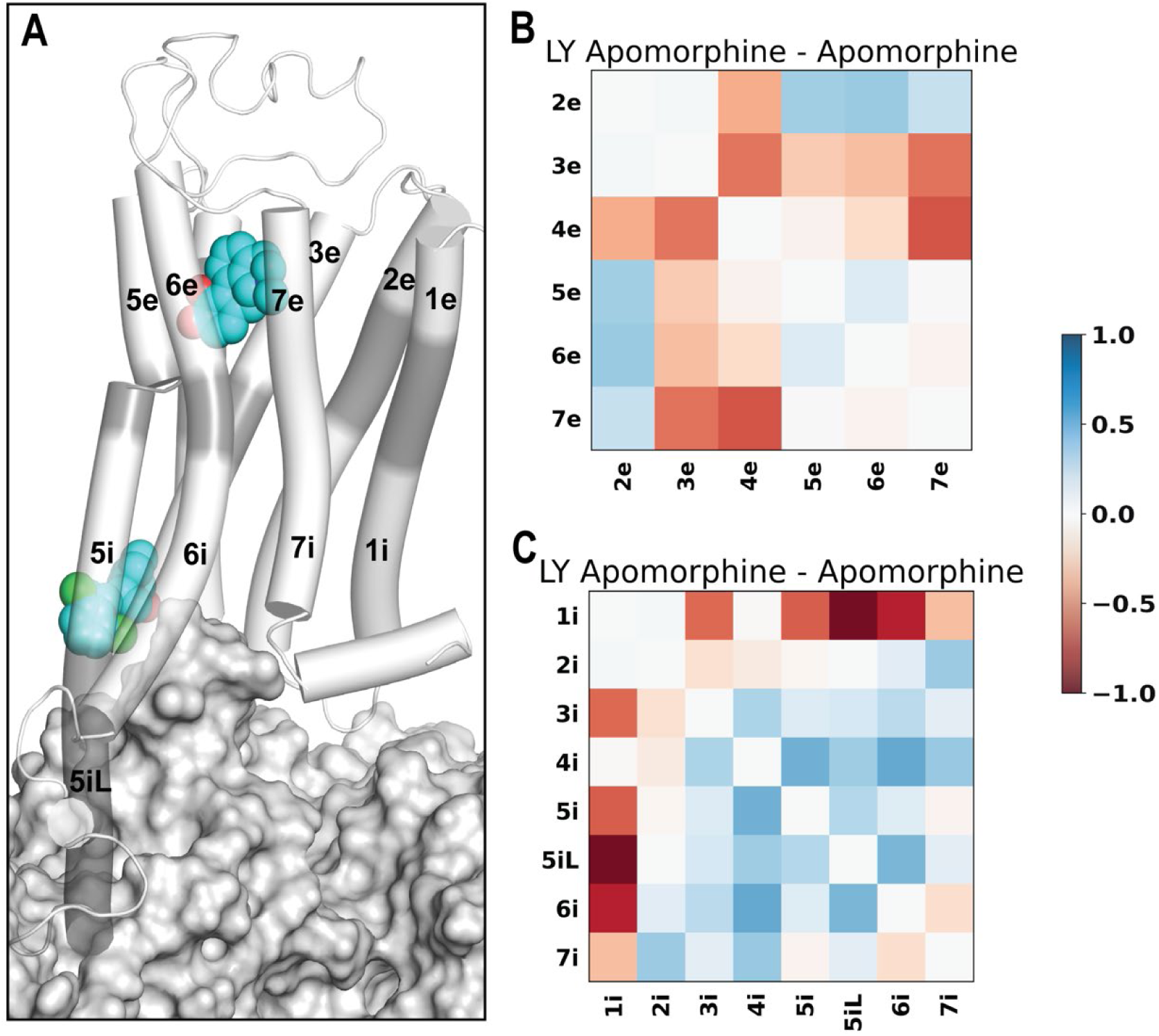
LY3154207 binding is associated with a tightening and loosening of the extracellular and intracellular regions of the receptor, respectively. (A) An overview of the D1R coupled with Gs protein and in complex with both apomorphine and LY3154207. The extracellular and intracellular subsegments of the D1R are colored in white, while the middle segments are in gray (see Methods for their definitions). Our PIA analysis demonstrates that the majority of extracellular subsegments have a tendency to move closer together (B), while the majority of intracellular subsegments have greater COM-COM distances (C) in the D1R/apomorphine-LY3154207 compared to the D1R/apomorphine-alone condition.

**Figure S7.**
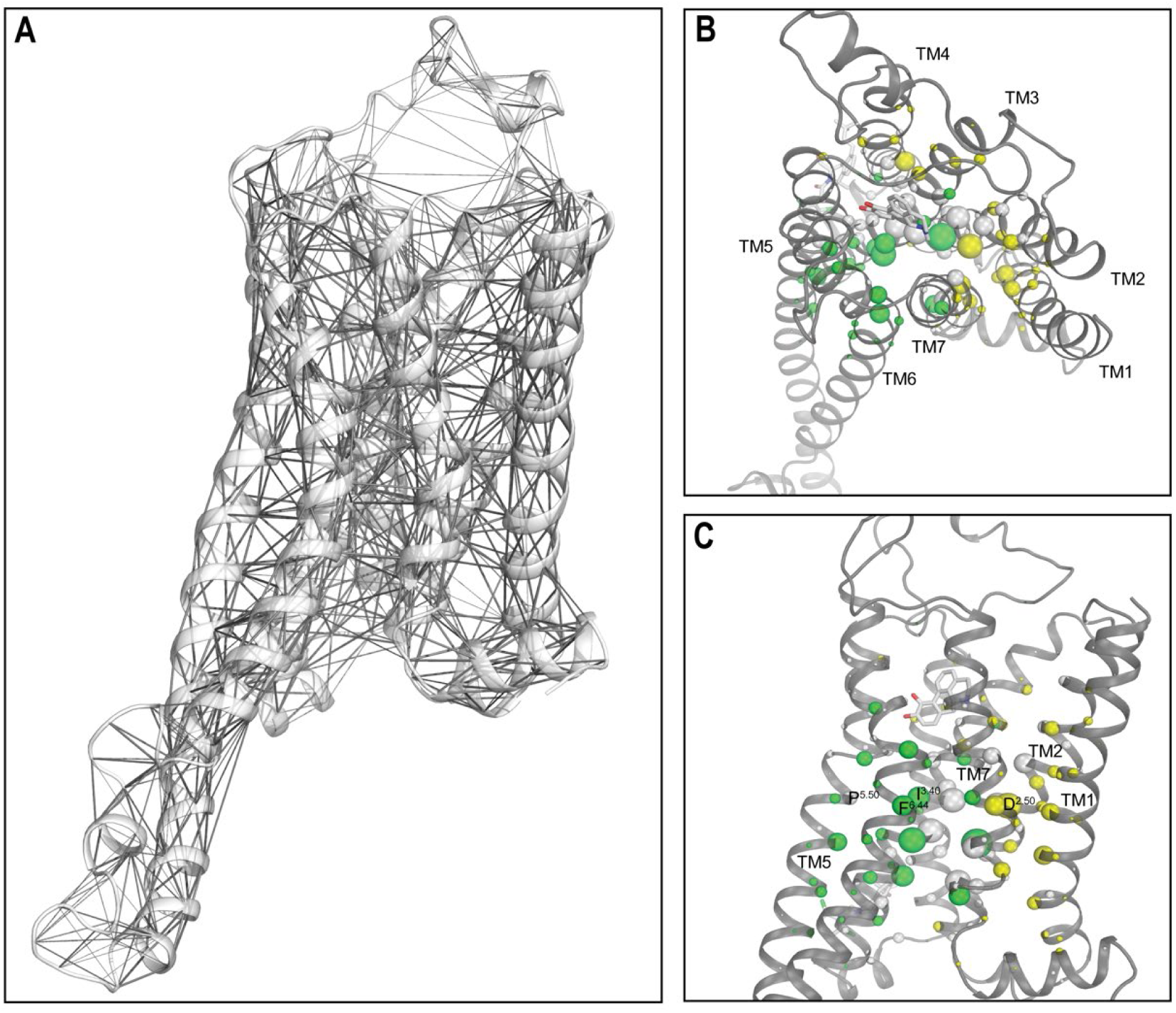
Network analysis identified impacts on several key structural motifs in the presence of LY3154207 in the D1R/apomorphine system. (A) A network connecting residues with larger than 0.2 contact frequency in the D1R/apomorphine-LY3154207 condition. If the shortest heavy-atom distance between two residues was within 4.5 Å, we defined that they formed a contact. Edge radii are scaled by contact frequency. (B) Top and (C) side views of the LY3154207-bound D1R/apomorphine system with nodes colored and scaled in radius according to the eigenvector centrality score. All nodes with an absolute score difference exceeding 0.005 are highlighted in color. Eigenvector centrality analysis demonstrates that in the presence of LY3154207, the D1R/apomorphine system has higher scores (yellow) for residues in the extracellular region enclosed by TMs 1, 2, 3, and 7, and lower scores (green) for residues in the intracellular region.

**Figure S8.**
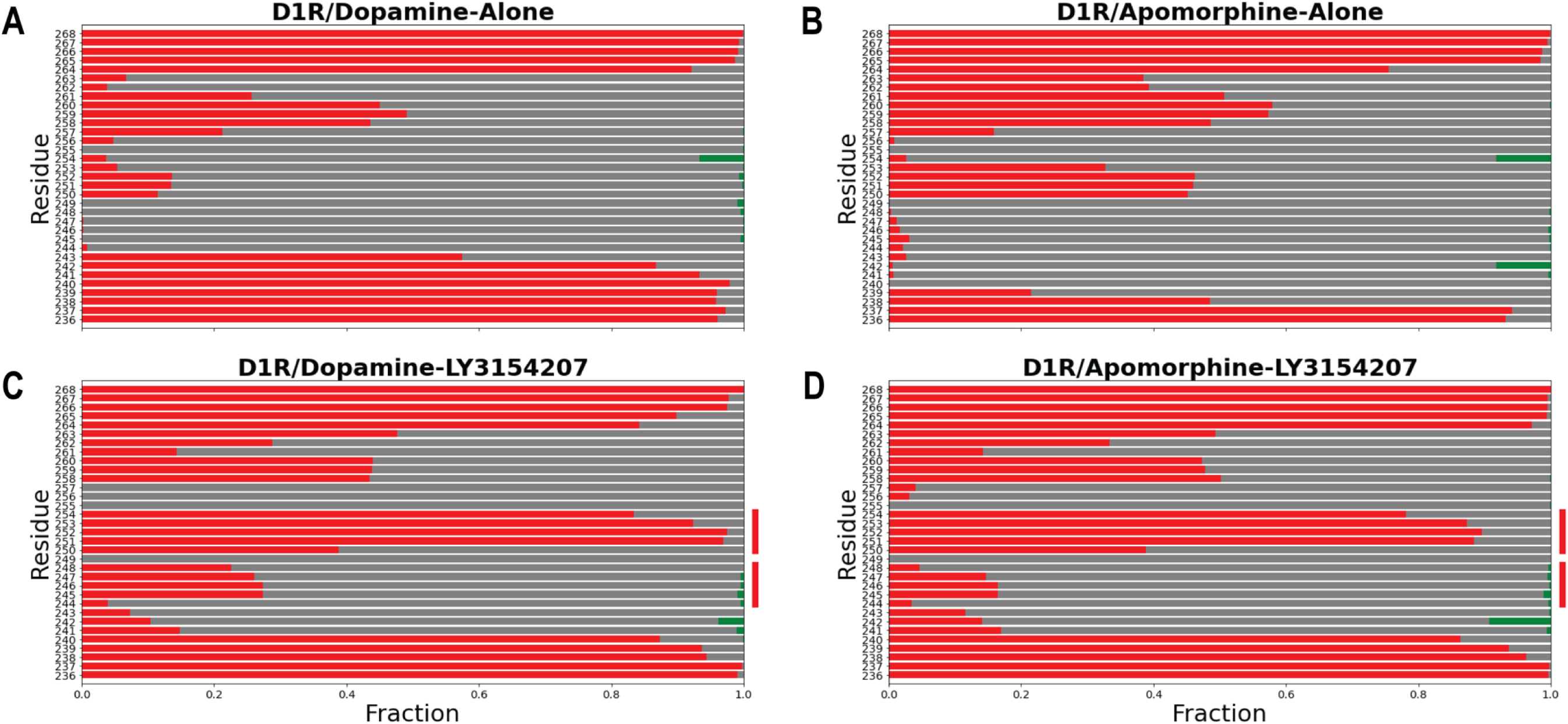
LY3154207 binding induces a conformational change in IL3. Secondary structure assignment for residues 236-268 of IL3. Helix, loop, and sheet assignment are depicted as red, gray, and green, respectively. In the D1R/dopamine (A) and D1R/apomorphine (B) systems, there is no convergent IL3 conformation. In the presence of LY3154207, the D1R/dopamine (C) and D1R/apomorphine (D) systems have similar IL3 conformations. Specifically, short helical segments emerge in IL3 for residues 245-248 and 250-254. Red bars are shown to the side of (C) and (D) to highlight regions that more frequently adopt a helical structure in the presence of LY3154207.

**Figure S9.**
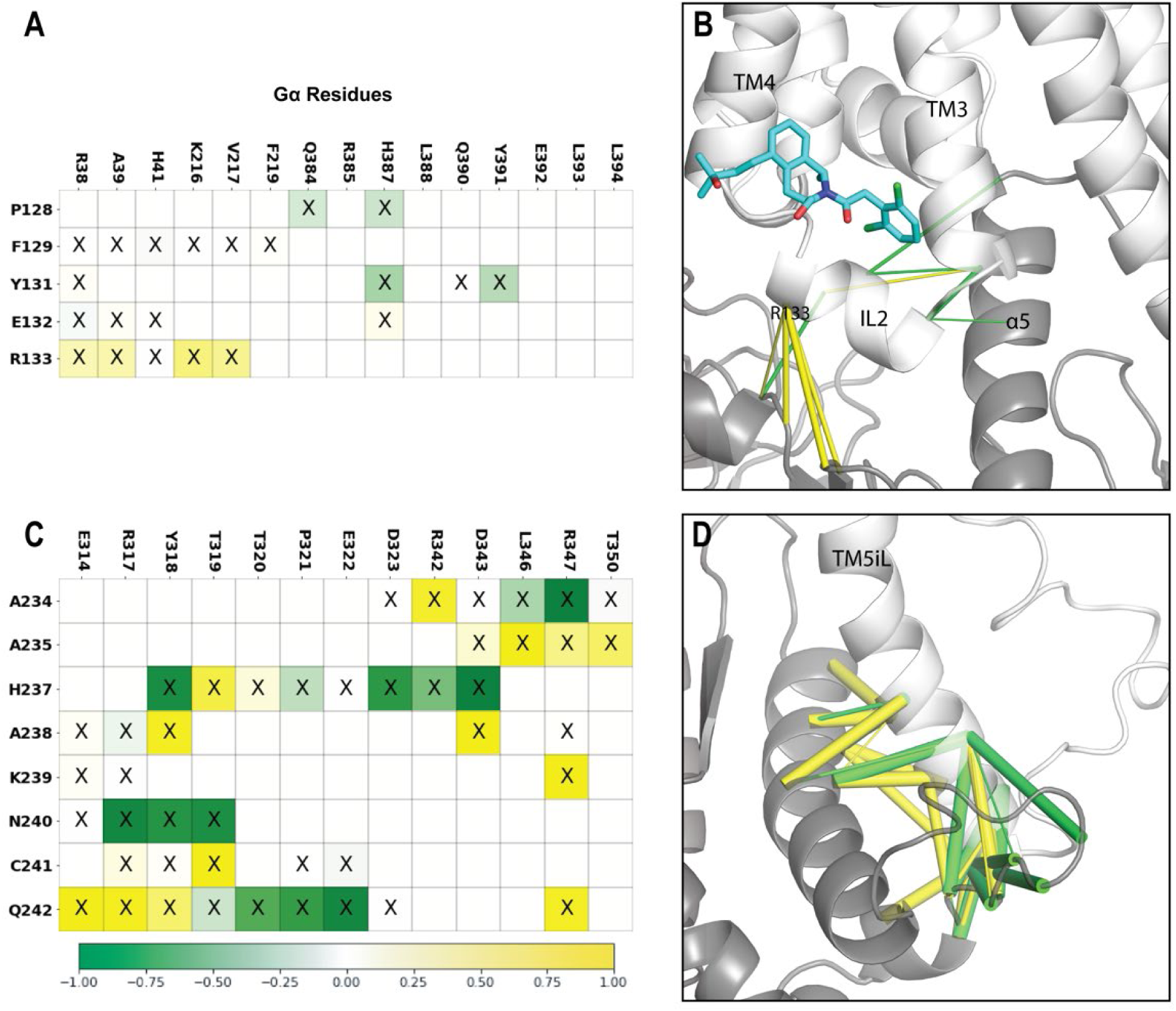
Contact frequency of receptor and G_sα_ residues in the D1R/apomorphine system. (A) The contact map for the interface between D1R/IL2 and Gα residues. Cells containing an ‘X’ represent the corresponding residue pairs forming a contact in both the D1R/apomorphine-LY3154207 and D1R/apomorphine-alone conditions. More frequent interactions in the presence of PAM are colored in yellow, and less frequent interactions are in green. Interactions are mapped on the D1R/Gα model shown in panel B. (B) In the presence of PAM, IL2 makes more consistent interactions with αN of Gα, specifically R133^IL2^. (C) The contact map for the interface between D1R/TM5iL and Gα residues is mapped to the model in panel (D). All contacts with a frequency difference greater than or equal to 0.10 are shown, and tubes are scaled to the frequency difference.

**Table S1.** Standardized histidine states. Histidine protonation states were standardized to those predicted by Protein Preparation Wizard using PROPKA pH 7.0.

| Chain | Residue | Protonation State |
| --- | --- | --- |
| **G<sub>α</sub>** | H41 | HID |
|  | H64 | HID |
|  | H220 | HID |
|  | H357 | HIP |
|  | H362 | HIP |
|  | H387 | HID |
| **G<sub>β</sub>** | H54 | HIE |
|  | H62 | HID |
|  | H91 | HID |
|  | H142 | HID |
|  | H183 | HID |
|  | H225 | HID |
|  | H266 | HID |
|  | H311 | HID |
| **G<sub>γ</sub>** | H44 | HID |
| **D1R** | H53 | HID |
|  | H164 | HID |
|  | H237 | HID |

**Table S2.**
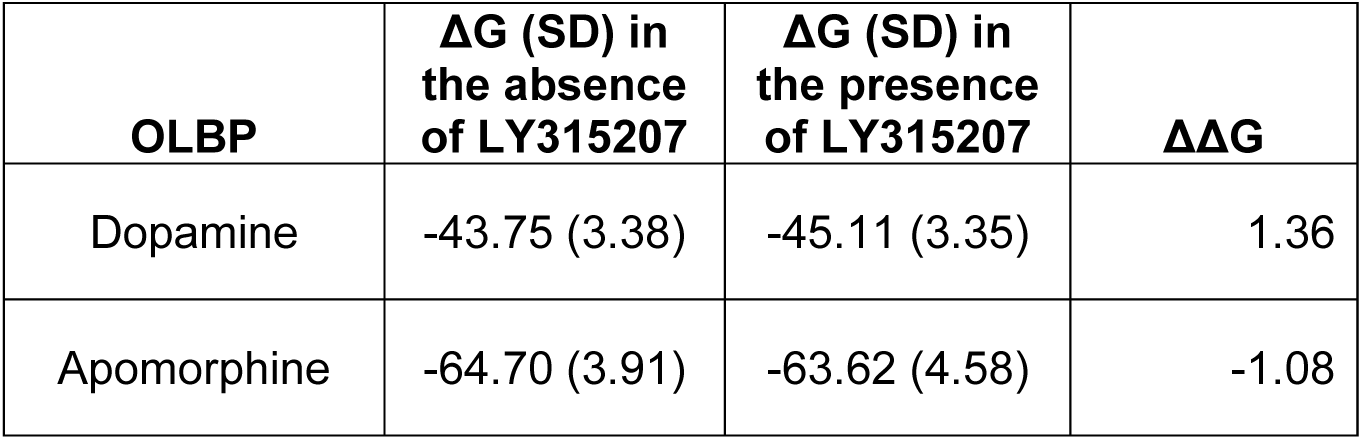
MM/GBSA calculations in the presence and absence of LY3154207. There is no consistent trend in the sign of ΔΔG values, however both systems show a ΔΔG below 1.4 kcal/mol, which are smaller than the standard deviations in each condition.

| **OLBP** | **$\Delta G$ (SD) in**<br>**the absence**<br>**of LY315207** | **$\Delta G$ (SD) in**<br>**the presence**<br>**of LY315207** | **$\Delta\Delta G$** |
| --- | --- | --- | --- |
| Dopamine | -43.75 (3.38) | -45.11 (3.35) | 1.36 |
| Apomorphine | -64.70 (3.91) | -63.62 (4.58) | -1.08 |

**Table S3.** The network eigenvector centrality scores of the D1R/apomorphine systems. Normalized eigenvector centrality scores for the D1R/apomorphine systems in the presence and absence of LY3154207. Only residues with an eigenvector centrality score of 0.10 or greater in any of the conditions are shown, and the score difference and absolute score difference are colored (green to yellow and gray to white) according to their respective maximum and minimum values.

| Residue | BW index | D1R/ApoM | D1R/ApoM-LY | ApoM - ApoM-LY | abs diff |
| --- | --- | --- | --- | --- | --- |
| 37 | TM1.46 | 0.075 | 0.118 | -0.043 | 0.043 |
| 41 | TM1.50 | 0.109 | 0.130 | -0.021 | 0.021 |
| 62 | TM2.42 | 0.171 | 0.173 | -0.002 | 0.002 |
| 63 | TM2.43 | 0.110 | 0.112 | -0.002 | 0.002 |
| 65 | TM2.45 | 0.118 | 0.119 | -0.001 | 0.001 |
| 66 | TM2.46 | 0.213 | 0.207 | 0.006 | 0.006 |
| 67 | TM2.47 | 0.087 | 0.095 | -0.008 | 0.008 |
| 69 | TM2.49 | 0.165 | 0.162 | 0.003 | 0.003 |
| 70 | TM2.50 | 0.158 | 0.176 | -0.018 | 0.018 |
| 72 | TM2.52 | 0.096 | 0.099 | -0.003 | 0.003 |
| 73 | TM2.53 | 0.139 | 0.140 | -0.001 | 0.001 |
| 77 | TM2.57 | 0.090 | 0.093 | -0.003 | 0.003 |
| 78 | TM2.58 | 0.086 | 0.102 | -0.016 | 0.016 |
| 105 | TM3.34 | 0.091 | 0.100 | -0.009 | 0.009 |
| 106 | TM3.35 | 0.120 | 0.119 | 0.001 | 0.001 |
| 107 | TM3.36 | 0.109 | 0.093 | 0.016 | 0.016 |
| 109 | TM3.38 | 0.121 | 0.119 | 0.002 | 0.002 |
| 110 | TM3.39 | 0.175 | 0.174 | 0.001 | 0.001 |
| 111 | TM3.40 | 0.155 | 0.147 | 0.008 | 0.008 |
| 113 | TM3.42 | 0.176 | 0.173 | 0.003 | 0.003 |
| 114 | TM3.43 | 0.203 | 0.183 | 0.020 | 0.020 |
| 117 | TM3.46 | 0.157 | 0.152 | 0.005 | 0.005 |
| 118 | TM3.47 | 0.109 | 0.097 | 0.012 | 0.012 |
| 148 | TM4.50 | 0.139 | 0.139 | 0.000 | 0.000 |
| 151 | TM4.53 | 0.099 | 0.100 | -0.001 | 0.001 |
| 203 | TM5.47 | 0.114 | 0.107 | 0.007 | 0.007 |
| 206 | TM5.50 | 0.105 | 0.100 | 0.005 | 0.005 |
| 210 | TM5.54 | 0.143 | 0.129 | 0.014 | 0.014 |
| 214 | TM5.58 | 0.101 | 0.082 | 0.019 | 0.019 |
| 281 | TM6.44 | 0.154 | 0.140 | 0.014 | 0.014 |
| 285 | TM6.48 | 0.127 | 0.122 | 0.005 | 0.005 |
| 321 | TM7.43 | 0.111 | 0.115 | -0.004 | 0.004 |
| 323 | TM7.45 | 0.114 | 0.104 | 0.010 | 0.010 |
| 324 | TM7.46 | 0.121 | 0.137 | -0.016 | 0.016 |
| 327 | TM7.49 | 0.154 | 0.152 | 0.002 | 0.002 |
| 328 | TM7.50 | 0.091 | 0.106 | -0.015 | 0.015 |
| 331 | TM7.53 | 0.155 | 0.138 | 0.017 | 0.017 |

## Notes

### Competing Interest Statement

The authors have declared no competing interest.

